# Genome-wide analysis of dendritic cell differentiation

**DOI:** 10.1101/2022.05.11.491577

**Authors:** Ioanna Tiniakou, Pei-Feng Hsu, Lorena S. Lopez-Zepeda, Colleen M. Lau, Chetna Soni, Eduardo Esteva, Nicholas M. Adams, Fan Liu, Alireza Khodadadi-Jamayran, Tori C. Rodrick, Drew Jones, Aristotelis Tsirigos, Uwe Ohler, Mark T. Bedford, Stephen D. Nimer, Boris Reizis

## Abstract

Dendritic cells (DCs) are immune sentinel cells that comprise antigen-presenting conventional DCs (cDCs) and cytokine-producing plasmacytoid DCs (pDCs). Cytokine Flt3 ligand (Flt3L) supports the proliferation of hematopoietic progenitors, and is also necessary and sufficient for DC differentiation. Here we characterized the spontaneous differentiation of a Flt3L-dependent murine progenitor cell line into pDCs and “myeloid” cDCs (cDC2s), and interrogated it using a genome-wide CRISPR/Cas9 dropout screen. The screen revealed multiple regulators of DC differentiation including the glycosylphosphatidylinositol transamidase complex, the Nieman-Pick type C cholesterol transporter and arginine methyltransferase Carm1; the role of Carm1 in pDC and cDC2 differentiation was confirmed by conditional targeting in vivo. We also found that negative regulators of mTOR signaling, including the subunits of TSC and GATOR1 complexes, restricted progenitor growth but enabled DC differentiation. The results provide a comprehensive forward genetic analysis of DC differentiation, and help explain how the opposing processes of proliferation and differentiation could be driven by the same cytokine.

## Introduction

Dendritic cells (DCs) link innate and adaptive immunity by recognizing pathogens through pattern recognition receptors such as TLRs and orchestrating antigen (Ag)- specific T cell responses (Steinman, 2012). DCs are represented by two main types: IFN-producing plasmacytoid DCs (pDCs) and Ag-presenting conventional DCs (cDCs). pDCs recognize pathogen-derived nucleic acids through endosomal TLRs (TLR7 and TLR9), and rapidly produce type I interferon (IFN) and other cytokines (Reizis, 2019; Swiecki and Colonna, 2015). cDCs have dendritic morphology, high levels of MHC class II and other components of Ag presentation machinery, express surface TLRs and can efficiently prime naive Ag-specific T cells. cDCs are comprised of two main subsets: XCR1^+^ cDC1s capable of antigen cross-presentation to CD8^+^ T cells and CD11b^+^ SIRPa^+^ cDC2s specialized in the presentation of exogenous antigen to CD4^+^ T cells (Guilliams et al., 2014).

DCs are short-lived mononuclear phagocytes that are being continuously produced in the bone marrow (BM), where they undergo subset specification followed by exit from the BM and, in the case of cDCs, by terminal differentiation in the periphery. DC development is governed by a hierarchy of transcription factors that specify the entire DC lineage (e.g. SPI1, IKZF1) or DC subsets including pDCs (e.g. TCF4, ZEB2) or cDC1 (e.g. IRF8, BATF3) (Murphy et al., 2016; Nutt and Chopin, 2020). The cell type-specific function of these factors correlates with their expression pattern, and is mediated via binding to specific gene targets. In contrast, little is known about other regulatory factors such as protein-modifying enzymes that may control DC development. Such putative enzymatic regulators are likely to be expressed ubiquitously and thus require unbiased genetic approaches for their identification.

The development of all DC subsets is driven by the cytokine Flt3 ligand (Flt3L). Its receptor Flt3 is expressed on and maintains the expansion of multipotent hematopoietic progenitors, but is also strongly expressed on committed DC progenitors and all mature DCs (Merad et al., 2013; Mildner and Jung, 2014). Both pDCs and cDCs, and only these cells, can develop in the sole presence of Flt3L from BM progenitors in vitro (Naik et al., 2005; Naik et al., 2007; Onai et al., 2007). All DCs are reduced in the lymphoid organs of Flt3L- or Flt3-deficient animals (McKenna et al., 2000; Waskow et al., 2008), whereas administration of Flt3L (Maraskovsky et al., 1996; Vollstedt et al., 2004) or its genetic activation through an oncogenic mutation (Lau et al., 2016) expands DC numbers. The activation of phosphatidylinositol-3-kinase (PI3K) signaling and mammalian target of rapamycin (mTOR) by Flt3 signaling is thought to be important for DC development (Sathaliyawala et al., 2010; Sukhbaatar et al., 2016; Weichhart et al., 2015). DC-specific deletion of Pten phosphatase, an upstream negative regulator of PI3K signaling, phenocopied Flt3L administration by causing an expansion of DC numbers in vivo (Sathaliyawala et al., 2010). Paradoxically, the deletion of TSC1, a negative regulator of mTOR, impaired DC differentiation in vitro and in vivo (Wang et al., 2013), suggesting that mTOR signaling may be detrimental to the process. Thus, the molecular mechanism of Flt3 signaling in DC development, and the role of mTOR signaling in the process, remain to be elucidated. This task requires a system that faithfully recapitulates both the Flt3L-dependent maintenance of DC progenitors and their differentiation into cDCs and pDCs.

Here we interrogated DC development in an unbiased manner by performing a CRISPR/Cas9-based dropout screen in a Flt3L-dependent DC progenitor cell line. The screen yielded multiple novel regulators including the arginine methyltransferase CARM1, whose specific role in DC development was confirmed *in vivo*. Importantly, the results revealed opposing roles of mTOR signaling in two discrete Flt3L-driven steps of DC development, i.e. progenitor proliferation and DC differentiation. The latter result identifies the inhibition of mTOR signaling as a critical molecular switch between proliferation and differentiation of hematopoietic progenitors driven by Flt3L.

## Results

### Modeling DC development in a Flt3L-dependent progenitor cell line

To model DC development in vitro, we used the Hoxb8-FL progenitor cells that are conditionally immortalized with the estrogen-inducible Hoxb8 oncogene and can be grown in the presence of estrogen and Flt3L as undifferentiated multipotent progenitors. Inactivation of Hoxb8 by estrogen removal in the presence of Flt3L induces a spontaneous program of DC differentiation that within 7 days yields a mixture of fully functional pDCs and cDCs (Bunin et al., 2015; Grajkowska et al., 2017; Redecke et al., 2013). While Hoxb8-FL cells have the potential to generate cDC1s once Notch signal has been provided, regular differentiation conditions yield only CD11b^+^ cDC2s (Kirkling et al., 2018). In our culture system, the majority of live cells on day 7 expressed DC marker CD11c and comprised SiglecH^+^ B220^+^ pDCs and CD11b^+^ MHC II^+^ cDC2s (Fig. S1A). First, we analyzed the transcriptome and open chromatin (by RNA-Seq and ATAC-Seq, respectively) of Hoxb8-FL-derived sorted pDCs and cDC2s. Representative RNA-Seq (Fig. 1A) and ATAC-Seq (Fig. 1B) sequence traces from Hoxb8-FL-derived DCs resembled those from primary DCs from the mouse spleen (Lau et al., 2018). Accordingly, global RNA-Seq and ATAC-Seq profiles of Hoxb8-FL-derived pDCs and cDC2s matched with their primary counterparts as assessed by hierarchical clustering (Fig. 1C,D) and principal component analysis (PCA) (Fig. S1B,C), after correcting for baseline differences between primary and in vitro conditions. Top variably expressed genes across all samples showed similar enrichment in either pDCs or cDCs irrespective of the cell source (Fig. S1D). Conversely, transcriptional and epigenetic signatures of primary splenic DC subsets (Lau et al., 2018) showed faithful expression in Hoxb8-FL-derived DCs: in particular, pan-cDC and cDC2 signatures were significantly enriched in cDC2s, and pDC signature was enriched in pDCs (Fig. 1E,F). In contrast, cDC1 signatures showed minor (RNA-Seq, Fig. 1E) or no (ATAC-Seq, Fig. 1F) enrichment in Hoxb8-FL-derived cDC2s, confirming that the latter largely correspond to cDC2s. Similarly, the transcripts for key pDC-specific transcription factors (*Tcf4, Irf8*) and markers (*Ly6D, Ccr9, Bst2, SiglecH*) were enriched in Hoxb8-FL-derived pDCs, whereas cDC2-enriched factors (*Notch2, Klf4*) and markers (*Sirpa, Itgam*) were enriched in Hoxb8-FL-derived cDC2s (Fig. 1G and S1E). Accordingly, Hoxb8-FL-derived cDC2s and pDCs showed the expected gene expression differences by pairwise comparison (Fig. S1F). The resolution of transcriptional (Fig. 1G) and epigenetic (Fig. 1H) subset signatures was less pronounced in Hoxb8-FL-derived DCs compared to primary DCs, likely reflecting the absence of relevant environmental signals such as lymphotoxin-beta and Notch ligands (Lewis et al., 2011). Nevertheless, these data suggest that Hoxb8-derived DC subsets represent adequate *in vitro* counterparts of primary pDCs and cDC2s.

**Figure 1.**
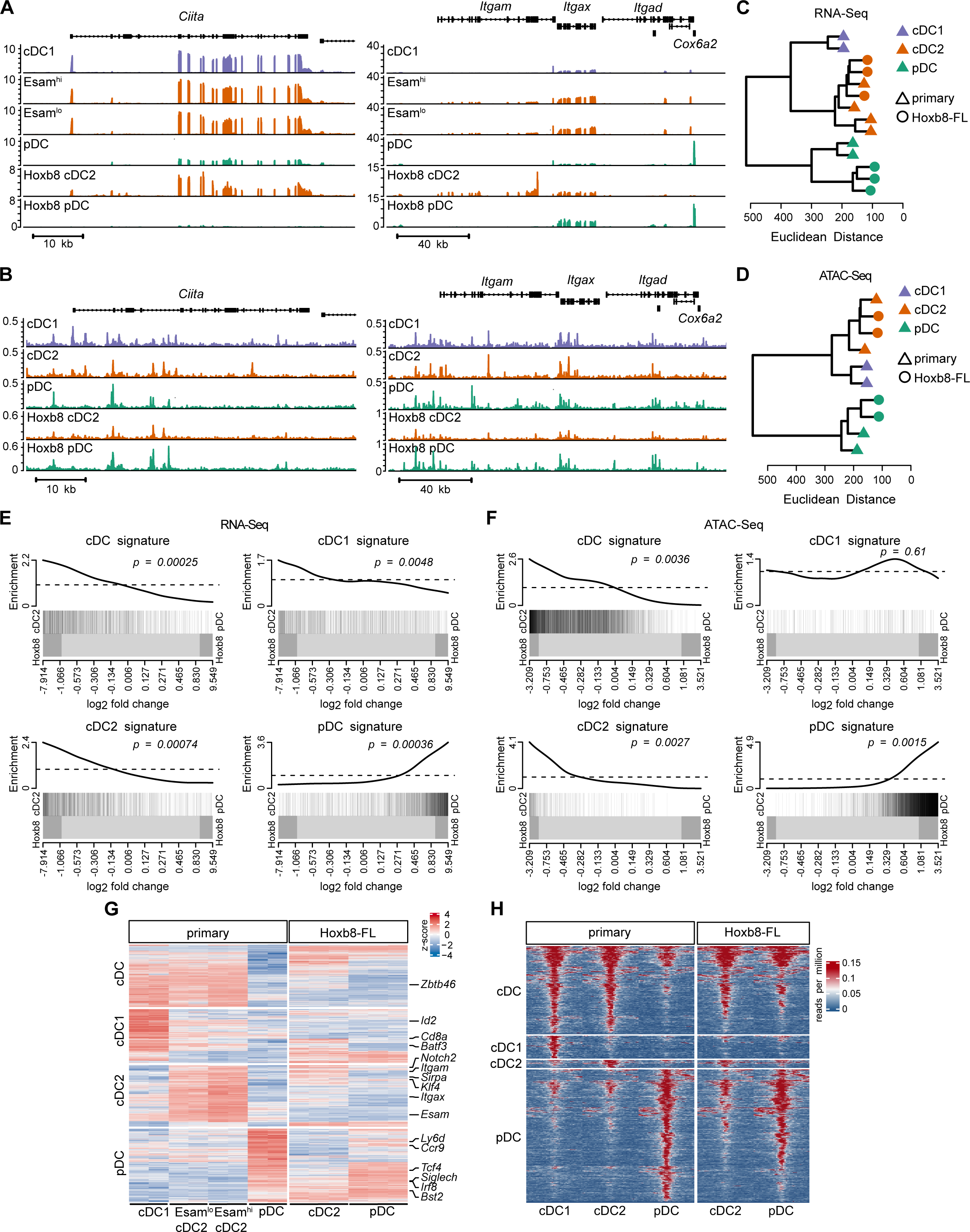
DCs derived from Hoxb8-FL cells resemble primary DCs in their transcriptome and epigenome. RNA-Seq and ATAC-Seq profiles of Hoxb8-FL-derived DCs were compared with those of primary splenic DCs from (Lau et al., 2018). (A-B) Representative RNA-Seq (panel A) and ATAC-Seq (panel B) profiles of DC-enriched genes in primary splenic DCs and Hoxb8-FL-derived DCs. Shown are normalized read counts across the indicated genomic loci. (C-D) Hierarchical clustering of global RNA-Seq (panel C) and ATAC-Seq (panel D) profiles of primary splenic DCs and Hoxb8-FL-derived DCs. Data are represented as dendrograms of Euclidean distances between samples calculated from log2 values and corrected for cell source (primary vs. Hoxb8-FL). (E-F) Enrichment of gene signatures of primary splenic DC subsets between Hoxb8-FL-derived DC subsets. Shown are barcode plots of transcriptomic signatures (RNA-Seq, panel E) and epigenomic signatures (ATAC-Seq, panel F) comparing Hoxb8-FL-derived cDC2s and pDCs and using log2 fold change gene expression values as ranked statistics. Black bars represent genes annotated with the gene set. Line shows relative enrichment of bars. Dark gray boxes indicate regions with absolute log2 fold change above 1. P-values are calculated from two-sided tests described in the Methods. (G) Transcriptomic gene signatures of DC subsets in primary splenic DCs and Hoxb8-FL-derived DCs. Shown is row-scaled heat map of log2-transformed gene expression values (rows) in biological replicates (columns), with select subset marker genes indicated. (H) Epigenomic signatures of DC subsets in primary splenic DCs and Hoxb8-FL-derived DCs. Shown is heatmap of peak-centered normalized counts from averaged ATAC-Seq data, with peaks plotted in 50-bp bins across a 2 kb window, and values beyond the 75th percentile ± 3 interquartile ranges capped.

Our recent lineage tracing-based analysis of DC differentiation in vivo (Feng et al., 2022) showed that both cDCs and pDCs emerge simultaneously from Cx3cr1^+^ progenitors, the earliest of which express Csf1R (CD115). More advanced DC progenitors (pro-DCs) upregulate CD11c and SiglecH, and subsequently committed pDC progenitors (pro-pDCs) downregulate Csf1R and Cx3cr1 and upregulate pDC markers Ly-6D and B220. We analyzed the process of Hoxb8-FL cells differentiation by staining them daily for markers of DCs and their progenitors. Differentiating Hoxb8-FL cells gradually downregulated the stem/progenitor marker CD34 and acquired DC marker CD11c by day 4 (Fig. 2A, top row). At that stage, they also acquired SiglecH and CD11b, which were subsequently resolved into SiglecH^+^ CD11b^-^ pDCs and SiglecH^lo^ CD11b^+^ cDC2s by day 7 (Fig. 2A, middle row). Notably, Csf1R and Cx3cr1 were rapidly induced on days 1-2 and were co-expressed on all cells by day 4, with a Csf1R^-^ population emerging on days 5-7 (Fig. 2A, bottom row). These Csf1R^-^ cells downregulated Cx3cr1 and acquired B220 and partially Ly-6D to yield pDCs, whereas the Csf1R^+^ population maintained Cx3cr1 and upregulated MHC II to become cDC2s (Fig. 2B). Thus, the differentiation of Hoxb8-FL cells closely recapitulates DC development *in vivo*, and includes the emergence of early Cx3cr1^+^ Csf1R^+^ progenitors on days 1-2, DC commitment by day 4 and the resolution of pDC and cDC2 subsets by day 7.

**Figure 2.**
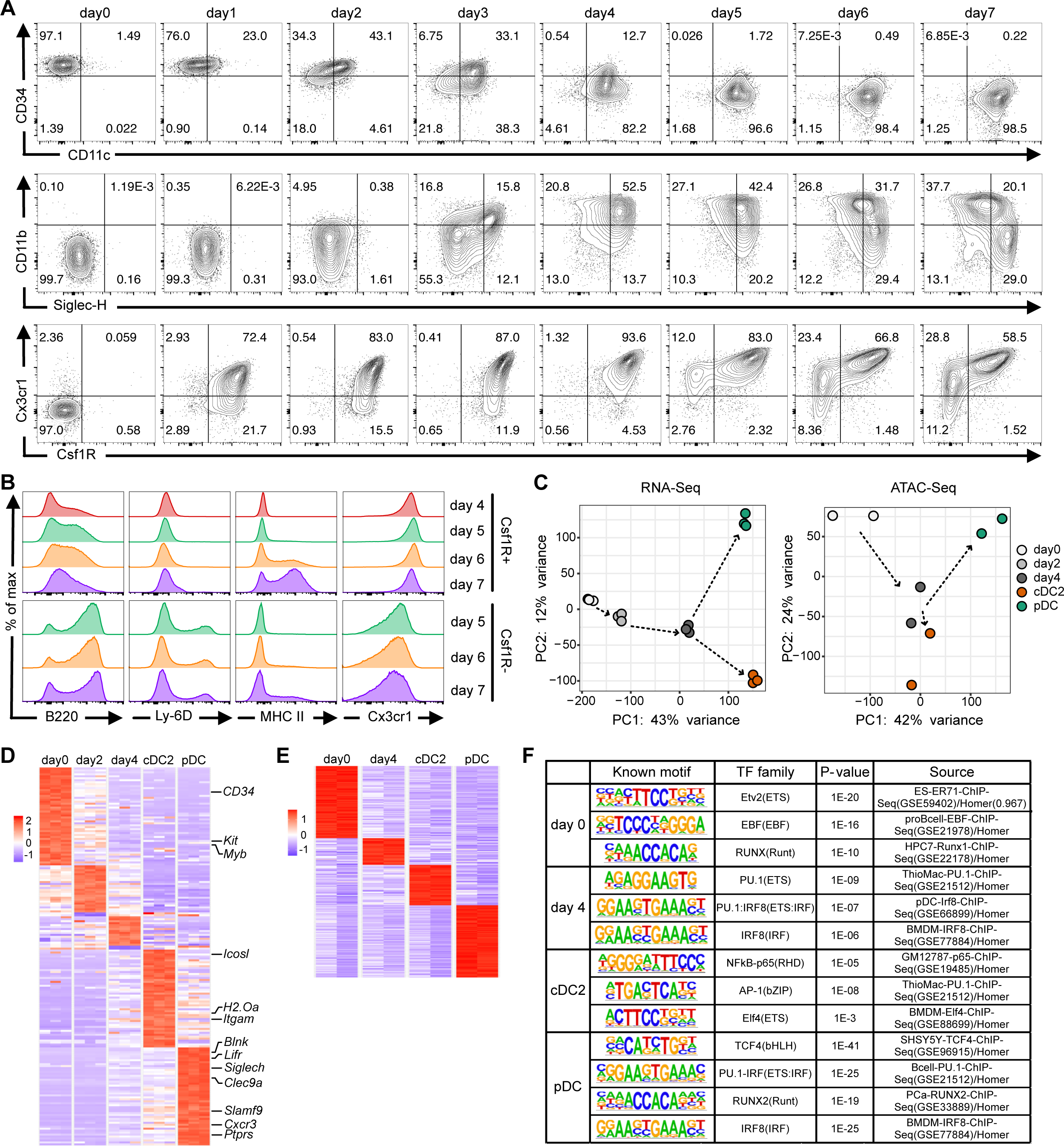
Differentiation of Hoxb8-FL progenitors recapitulates DC differentiation in vivo. (A) Surface marker expression in live Hoxb8-FL cells on days 0-7 of differentiation. (B) Expression of indicated surface markers on gated CSF1R^+^ or CSF1R^-^ cells on days 4-7 of differentiation. (C) Principal component analysis of global RNA-Seq (left panel) or ATAC-Seq (right panel) profiles of Hoxb8-FL cells at the indicated days of differentiation (0-4) and in differentiated cDC2s and pDCs (day 7). Symbols represent biological replicates; temporal relationships between samples are indicated by dashed arrows. (D-E) Stage-specific signatures of Hoxb8-FL cell differentiation. Heatmaps show row- scaled log2-transformed gene expression values (RNA-Seq, panel D) or peak values (ATAC-Seq, panel E) in biological replicates (columns) of Hoxb8-FL cells at the indicated days of differentiation. Select subset-enriched marker genes are indicated. (F) Transcription factor (TF) binding motif enrichment in epigenomic signatures of differentiation stages shown in panel E. Shown are the results of HOMER motif enrichment analysis with the known motif, top TF match and TF family, p-value of the match and the data source of the top match.

Having established the kinetics of Hoxb8-FL differentiation, we extended their transcriptional and epigenetic profiling to undifferentiated Hoxb8-FL progenitors (day 0) and committed DC progenitors prior to subset specification (day 4). An additional stage of early DC progenitors (day 2) was used for RNA-Seq only. PCA of the resulting RNA-Seq profiles confirmed stepwise differentiation of these populations towards DCs (PC1) and resolution of DC subsets on day 7 (PC2) (Fig. 2C). Accordingly, DC-enriched transcripts were upregulated on day 4 and were resolved into common or subset-specific sets on day 7 (Fig. S2A). PCA of ATAC-Seq profiles suggested a similarity between committed DC progenitors (day 4) and cDC2s, whereas pDCs were further resolved from the latter along both PC1 and PC2 (Fig. 2C). Accordingly, both pDCs and cDC2s showed major differences from day 4 cells by RNA-Seq (Fig. S2B), whereas ATAC-Seq revealed profound changes only in pDCs (Fig. S2C). As an example, RNA-Seq showed gradual upregulation of key transcription factors such as *Tcf4* and *Irf8* in pDCs and *Id2* in cDC2s (Fig. S2D). At the same time, the associated chromatin features were observed primarily in pDCs, such as pDC-specific enhancers of *Tcf4* (Grajkowska et al., 2017) and *Irf8* (Bagadia et al., 2019) and the negative distal regulatory region of *Id2* (Ghosh et al., 2014) (Fig. S2E). We then defined sets of genes (for RNA-Seq) and peaks (for ATAC-Seq) specific for each differentiation stage (Fig. 2D,E). pDC-specific ATAC-Seq peaks were more abundant than those specific for day 4 or cDC2s (Fig. 2E), further suggesting that pDC differentiation of Hoxb8-FL cells is associated with deeper epigenetic reprogramming. The analysis of transcription factor binding motifs enriched in stage-specific peak sets (Fig. 2F) showed that day 4 was enriched in PU.1, IRF8 and combined PU.1:IRF8 motifs, reflecting the requirement for these factors in early DC development (Carotta et al., 2010; Murakami et al., 2021; Sichien et al., 2016). Mature pDCs showed enrichment of motifs for TCF4 and RUNX2, two factors critical for pDC development and function (Cisse et al., 2008; Sawai et al., 2013). Conversely, cDC2s showed the expected lack of enrichment for IRF8 binding motifs but an enrichment of motifs for NF-κB, which is required for cDC development (Ouaaz et al., 2002). Collectively, these results provide a comprehensive molecular description of stepwise DC differentiation and support Hoxb8- FL cells as its relevant model.

### Genome-wide genetic analysis of Hoxb8-FL cell growth and differentiation

To characterize Flt3L-dependent growth and differentiation of Hoxb8-FL cells, we performed a CRISPR/Cas9-based dropout screen using a genome-wide library of single guide RNAs (sgRNAs). We generated a Cas9-expressing Hoxb8-FL clone with an intact differentiation capacity (Fig. S3A,B) and then transduced it with the “Brie” sgRNA library targeting ∼19,600 mouse genes with 4 sgRNA per gene (Doench et al., 2016). Cells were transduced at MOI<0.5 to limit multiple integrations, resulting in an estimated ∼3.2 x10^7^ primary transfectants (∼400 cells per sgRNA) (Fig. 3A). Cells growing in an undifferentiated state (in the simultaneous presence of Flt3L and estrogen) were selected with puromycin and expanded to yield the initial progenitor sample (progenitors_0). Additional expansion for 6 days yielded the sample of undifferentiated progenitors. The cells were then differentiated in Flt3L without estrogen for 7 days, and both total differentiated DCs and sorted pDC and cDC2 samples were collected. The library from each sample, as well as from the original plasmid DNA (pDNA) was amplified, sequenced and analyzed for sgRNA content. The distribution of targeting sgRNAs was progressively skewed in the progenitor and DC samples as reflected by reduced diversity index (Fig. S3C) and broader distribution of counts (Fig. S3D). The effect was much less pronounced in ∼1,000 non-targeting sgRNAs within the library, suggesting that it reflects functional changes.

**Figure 3.**
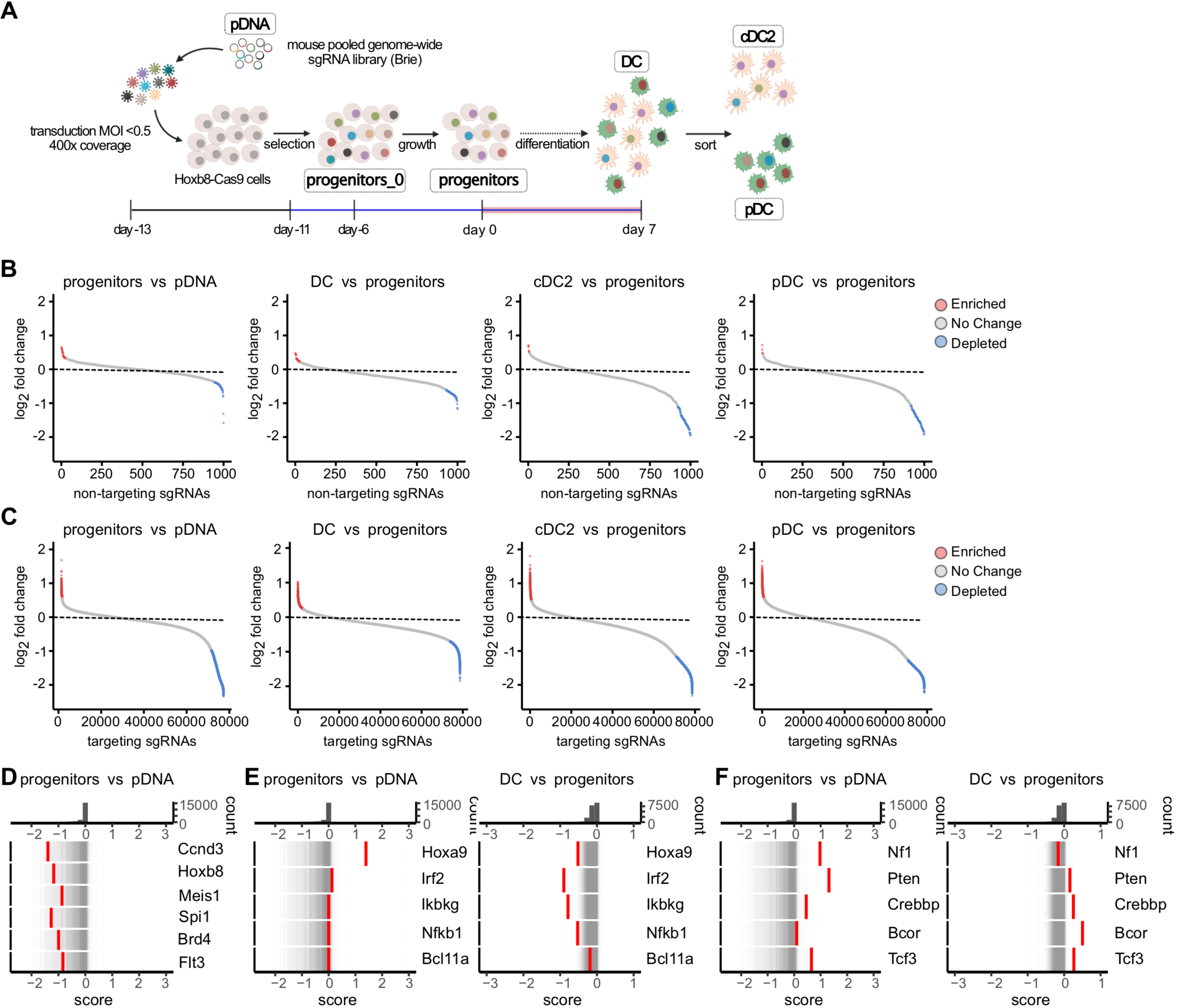
Genome-wide screen for regulators of Hoxb8-FL cell proliferation and differentiation. (A). Schematic of the screen, indicating the samples collected for sgRNA library analysis. (B-C). Pairwise comparison of individual sgRNA levels in collected samples. Shown are ranked log2 fold changes in the abundances of individual non-targeting control sgRNAs (panel B) and targeting sgRNAs (panel C) in the indicated sample pairs. Significantly enriched or depleted sgRNAs with false discovery rate (FDR) <0.05 are highlighted in red and blue, respectively. (D-F). Rug plots of gene distribution between indicated pairs of samples as measured by the CRISPTimeR score. Top panels show the distribution histogram of scores; bottom panels show individual genes (red bars) overlaid on the distribution of all genes (thin grey bars). (D) Rug plots of select genes showing sgRNA depletion in progenitors. (E) Rug plots of select genes showing sgRNA depletion in DCs. (F) Rug plots of select genes showing sgRNA enrichment in progenitors and/or DCs.

The comparison of progenitor sample to pDNA should identify genes that regulate the growth of undifferentiated progenitors. Accordingly, 18.2% of targeting sgRNAs were depleted in progenitors vs pDNA, compared to 7.1% of non-targeting sgRNAs (Fig. 3B,C, Fig. S3E). The comparison of DC to progenitor samples should identify genes that regulate DC differentiation. Indeed, 32.9% of targeting sgRNAs vs 19.1% of non-targeting sgRNAs were depleted in the total DC sample (Fig. 3B,C and Fig. S3E). The difference between targeting and non-targeting sgRNAs was lower but still detectable in sorted pDCs and cDC2s, likely reflecting the lower cell numbers in these samples. The fractions of targeting sgRNAs that were enriched in progenitors or in DCs were similar to those of non-targeting RNAs (Fig. S3E), suggesting that the screen predominantly detects sgRNA depletion in both cases.

The samples were then compared using the CRISPTimeR algorithm that identifies hits in dropout screens (Garipler et al., 2022). Depletion of multiple sgRNAs for a given gene yields a low CRISPTimeR score and identifies this gene as a positive regulator; conversely, sgRNA enrichment produces a high CRISPTimeR score corresponding to a negative regulator. As expected, genes with low CRISPTimeR score in progenitors were enriched for genes involved in essential biological processes such as biosynthesis, translation and ribosome biogenesis (Fig. S3F). Indeed, the top 1,000 low-score genes in progenitors included 123/217 (57%) of “gold-standard” human essential genes but only 1/927 (0.1%) of non-essential genes (Morgens et al., 2016). Notably, they also included the lymphoid progenitor-specific cyclin *Ccnd3* (Sicinska et al., 2003), the transforming oncogene *Hoxb8*, the essential growth factor receptor *Flt3*, and key transcriptional regulators of hematopoietic/DC progenitors such as *Meis1*, *Spi1* (encoding PU.1) and *Brd4* (Fig. 3D). On the other hand, genes with neutral scores in progenitors vs pDNA but low scores in DCs vs progenitors included known transcriptional regulators of DC differentiation: *Irf2*, which promotes cDC2 development by restricting IFN-I signaling (Honda et al., 2004; Ichikawa et al., 2004); *Nfkb1* and *Ikbkg*, components of the NF-κB pathway that controls DC differentiation (Ouaaz et al., 2002); *HoxA9*, which facilitates Flt3L-driven DC development *in vitro* (Bleyl, 2018); and *Bcl11a*, which is required in committed DC progenitors and pDCs (Ippolito et al., 2014; Wu et al., 2013) (Fig. 3E). These genes had negative scores in both pDC and cDC2 samples (Fig. S3G), suggesting the activity in committed progenitors prior to subset specification; nevertheless, *Irf2* and *Bcl11a* showed more profound depletion in cDC2s and pDCs, respectively, consistent with their requirements in these subsets.

Among the few genes with higher CRISPTimeR score (i.e. negative regulators) was *Pten*, consistent with its negative role in cDC differentiation (Sathaliyawala et al., 2010). *Pten* showed positive score in both progenitors and DCs, in contrast to other tumor suppressors such as *Nf1* that was enriched only in progenitors (Fig. 3F). Other high-score genes in both progenitors and DCs were *Crebbp*, encoding the transcriptional coactivator CBP; and *Tcf3*, encoding the lymphoid transcription factor E2A. The latter effect is not consistent with the proposed role of E2A in pDC development (Bagadia et al., 2019) and requires further study. Notably, the top gene with high score specifically in DCs was *Bcor*, encoding a transcriptional corepressor that was recently identified as a negative regulator of Flt3L-driven DC differentiation (Tian et al., 2021). The enrichment of these genes was detectable in cDC2s but less pronounced in pDCs (Fig. S3H), likely due to technical limitations of this sample. Thus, the screen was able to identify multiple known regulators of two separate Flt3L-driven processes, i.e. progenitor proliferation and DC differentiation.

### Identification of additional regulators of DC differentiation

We explored the ability of our screen to identify additional regulators of DC differentiation. The genes with the lowest CRISPTimeR scores in DCs (but normal scores in progenitors) included all five subunits of the glycosylphosphatidylinositol (GPI) transamidase complex (*Pigc, Pigk, Pigt, Pigu* and *Gpaa1*), which adds GPIs to newly synthesized proteins (Yu et al., 2013) (Fig. 4A). Additional low-scoring genes included *Setdb1*, a histone lysine methyltransferase required for hematopoietic cell differentiation (Koide et al., 2016); *Jmjd6*, a JmjC domain-containing enzyme with multiple critical functions (Kwok et al., 2017); *Ppp2ca*, a catalytic subunit of Ser/Thr phosphatase 2A; and several others including *Morc3*, *Colgalt1* and *Ubr4* (Fig. 4A). All these genes had low scores in both pDCs and cDC2s, suggesting their general role in DC differentiation (Fig. 4A). To confirm the role of these genes in DC differentiation, they were targeted by individual sgRNAs in Hoxb8-FL cells. Two sgRNAs per gene were used (one from the original Brie library and one designed independently) and showed similar results. All resulting batch cultures showed normal growth (data not shown) but defective DC differentiation, ranging from a near-complete block (GPI transamidase subunits, *Ppp2ca*) to a partial impairment (*Setdb1*, *Jmjd6*) (Fig. 4B).

**Figure 4.**
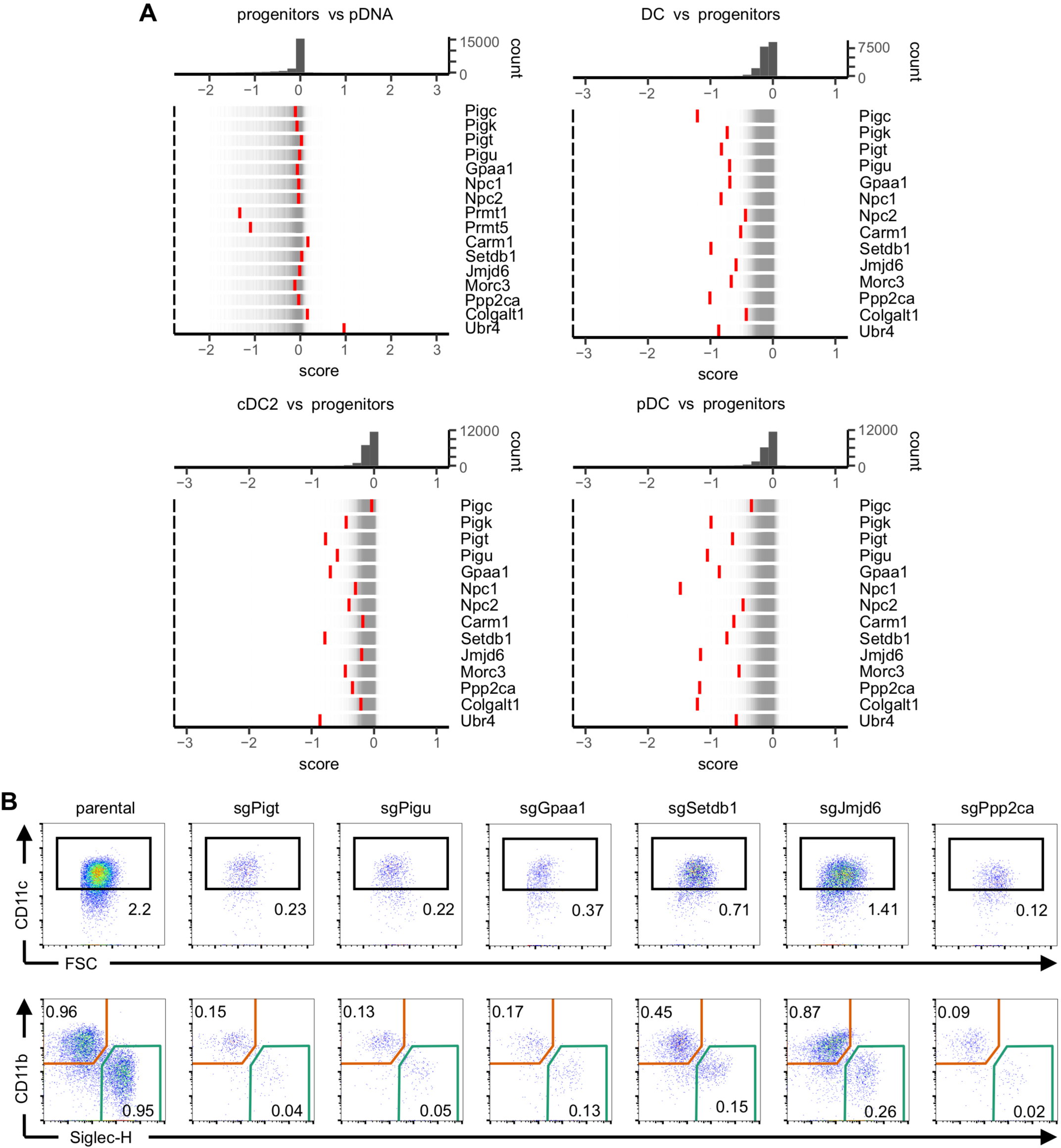
Identification of candidate regulators of DC differentiation. (A) Rug plots of gene distribution between indicated pairs of samples, as measured by the CRISPTimeR score. Top panels show the distribution of scores; bottom panels show select candidate genes (red) against the distribution of all genes (grey). (B) Differentiation capacity of Hoxb8-FL cells deficient for select candidate genes from panel A. Parental Hoxb8-FL cells or cells transduced with sgRNAs targeting indicated genes were differentiated into DCs and analyzed by flow cytometry as in Fig. S3B. Upper row shows gated viable singlets with the CD11c^+^ DCs highlighted; lower row shows gated CD11c^+^ DCs with SiglecH^+^ pDCs and CD11b^+^ cDC2s highlighted. Numbers indicate the fraction of cells in the gate among total singlets. Representative of two sgRNAs per gene.

Genes with the lowest score in DCs (Fig. 4A) included *Npc1* and *Npc2*, which encode two subunits of the NPC complex that exports cholesterol from lysosomes to cytosol (Pfeffer, 2019; Subramanian and Balch, 2008) Mutations in either of these genes cause Niemann-Pick type C disease characterized by intracellular cholesterol accumulation and age-dependent fatal neurotoxicity, which is recapitulated in Npc1- deficient mice. We analyzed these *Npc1*^-/-^ mice prior to the neurotoxicity phenotype and found normal cellularity of the BM and lymphoid organs (Fig. S4A) and normal fractions of myeloid cells (Fig. S4B,C) compared to littermate *Npc1*^+/+^ controls. However, the fraction of pDCs was significantly reduced in the BM and showed a similar trend (albeit not reaching significance) in the spleen (Fig. S4D,E). Moreover, the fraction of splenic cDCs was significantly reduced due to a specific ∼2-fold reduction of cDC2s but not cDC1s (Fig. S4F). We then transferred BM from *Npc1*^-/-^ or control *Npc1*^+/+^ mice mixed with wild-type CD45.1^+^ competitor BM into irradiated CD45.1^+^ recipient mice. In the resulting mixed chimeras, donor-derived pDCs were underrepresented ∼2-fold in the recipients of *Npc1*^-/-^ BM, although they were predominant over competitor-derived pDCs in recipients of *Npc1*^+/+^ BM (Fig. S4G). Similarly, the fraction of donor-derived splenic cDC2s was reduced in *Npc1*^-/-^ BM recipients, although the results did not reach statistical significance due to limited numbers (Fig. S4H). These results demonstrate the role of *Npc1* in DC development in vivo, and suggest that it reflects a cell-intrinsic activity of the NPC complex. Collectively, our data support the ability of our unbiased screening platform to identify functional regulators of DC differentiation.

### Carm1 regulates DC differentiation in vitro and in vivo

Methylation of arginine residues is a critical protein modification that controls multiple aspects of cellular physiology, and is mediated by a family of protein methyltransferase (PRMT) enzymes (Guccione and Richard, 2019). Indeed, genes for constitutive PRMT enzymes, *Prmt1* and *Prmt5,* showed very low scores in Hoxb8-FL progenitors, suggesting their essential role in cell growth (Fig. 4A). In contrast, *Prmt4* (also known as coactivator-associated arginine methyltransferase 1 or *Carm1*) was unaffected in progenitors but showed low score in DCs, particularly in pDCs (Fig. 4A). To confirm the specific role of *Carm1* in DC differentiation, we targeted it individually in Hoxb8-FL cells using two different sgRNAs, both of which led to the loss of CARM1 protein and of its specific substrate, arginine-methylated PABP1 (Lee and Bedford, 2002) (Fig. 5A). The resulting Carm1-deficient Hoxb8-FL cells showed normal growth (data not shown) but impaired differentiation into DCs, particularly into pDCs (Fig. 5B), whereas the few resulting cDCs had low MHC cl. II levels (Fig. S5A). Accordingly, small molecule inhibitors of CARM1 specifically impaired pDC differentiation from Hoxb8-FL cells (Fig. S5B), further supporting the role of CARM1 in this process.

**Figure 5.**
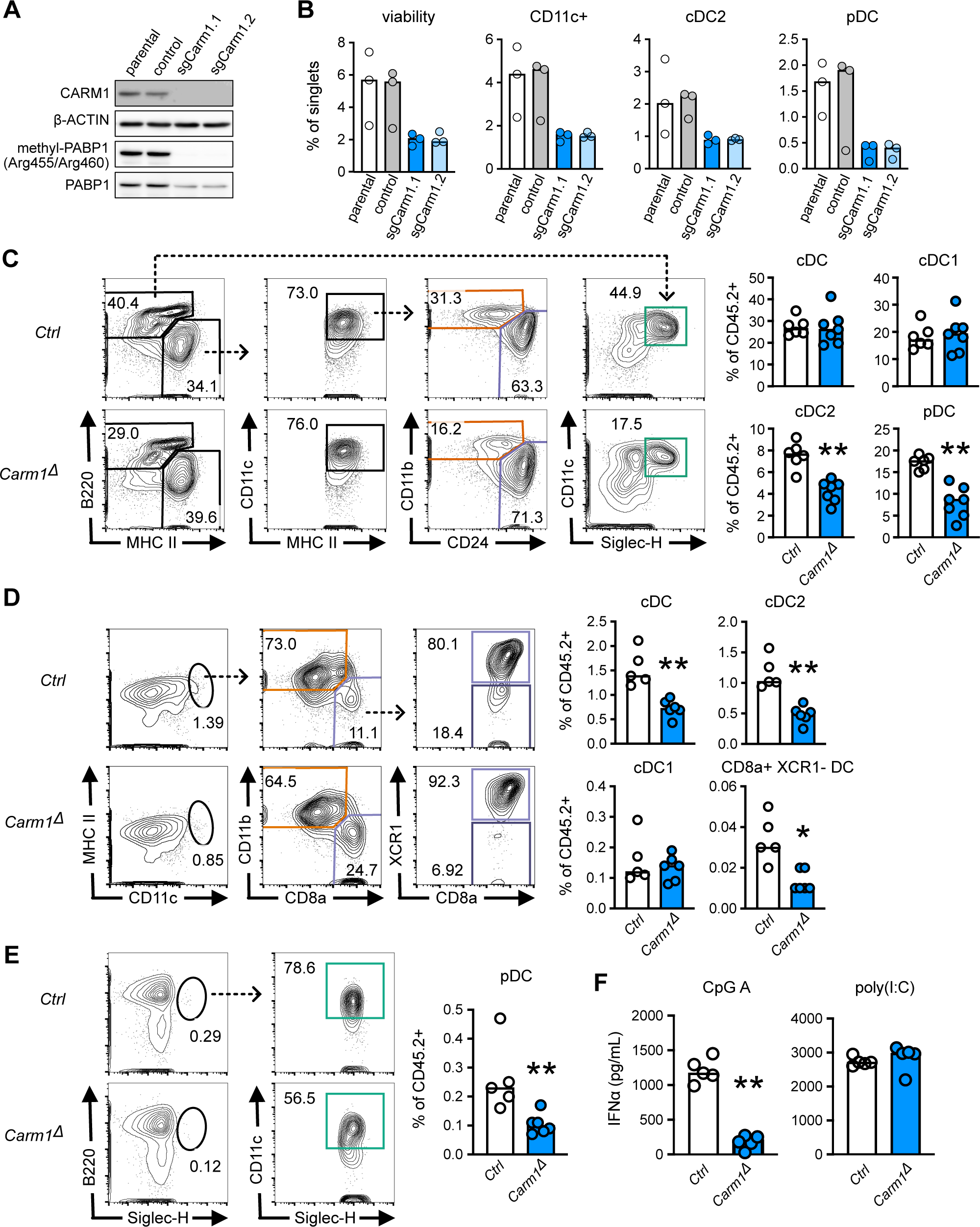
*Carm1* facilitates the differentiation of pDCs and cDC2s. (A) Deletion of *Carm1* in Hoxb8-FL cells. Cas9-expressing parental Hoxb8-FL cells were transduced with lentiviruses encoding control sgRNA or sgRNAs against *Carm1* (sgCarm1.1 or sgCarm1.2). Shown is Western blot of undifferentiated cells probed for CARM1, its substrate PABP1 methylated on Arg455 and Arg460, total PABP1 or β-ACTIN as a loading control. (B) Differentiation of *Carm1*-deficient Hoxb8-FL cells. Parental Hoxb8-FL cells or cells transduced with control or *Carm1*-targeting sgRNAs (panel A) were differentiated into DCs and analyzed by flow cytometry as in Fig. S3B. Shown are the fractions of viable cells, total CD11c^+^ DCs or DC subsets among single cells. Symbols represent separate experiments; bars represent the median. (C-F) Ablation of *Carm1* in hematopoietic cells *in vivo*. CD45.1^+^ recipient mice were reconstituted with BM from mice with a pan-hematopoietic deletion of *Carm1* (*Carm1*^fl/fl^ *Vav1*-Cre, designated *Carm1*^Δ^) or control (*Carm1*^fl/fl^) CD45.2^+^ mice, and analyzed eight months later. Donor reconstitution was >84% at one month post transplantation; all data reflect gating on CD45.2^+^ donor cells. (C) DC development in Flt3L-supplemented BM cultures. Shown are representative staining profiles of day 7 cultures, highlighting CD11c^+^ MHCII^+^ cDCs, CD11b^hi^ cDC2s, CD24^hi^ cDC1-like cells and SiglecH^+^ pDCs. Bar graphs show frequencies of indicated DC subsets among donor-derived cells. Symbols represent cultures from individual recipient mice (n = 6-7); bars represent the median. (D-E) Splenic cDCs (panel D) and pDCs (panel E) in recipient mice. Shown are representative staining profiles of CD45.2^+^ donor splenocytes, highlighting CD11c^+^ MHCII^+^ cDCs, CD11b^hi^ cDC2s, XCR1^+^ cDC1s, XCR1^-^ CD8a^+^ tDCs and SiglecH^+^ pDCs. Bar graphs show frequencies of indicated DC subsets among donor-derived splenocytes. Symbols represent individual recipient mice (n= 5-6); bars represent the median. (F) Production of IFN-I by recipient mice. Mice were injected i.v. with CpG-A and IFN-α concentration was measured in the serum 6 hours later. Three weeks later, the same animals were injected i.p. with poly(I:C) and serum IFN-α was measured 6 hours later by ELISA. Symbols represent individual recipient mice (n= 5); bars represent the median. Statistical significance was deterimined by Mann-Whitney test (*, P < 0.05; **, P < 0.01).

*Carm1*-deficient mice die of respiratory distress at birth (Yadav et al., 2003) but show normal hematopoiesis except for aberrant thymocyte development (Kim et al., 2004). Moreover, conditional targeting of *Carm1* showed that it is generally dispensable for hematopoietic cell differentiation in the adult (Greenblatt et al., 2018). To test the role of *Carm1* in DC development in vivo, we transferred BM from mice with a constitutive deletion of *Carm1* in hematopoietic cells (*Carm1*^fl/fl^ *Vav1*-Cre, designated *Carm1*^Δ^) or control *Carm1*^fl/fl^ mice (CD45.2^+^) into CD45.1^+^ congenic recipients. Consistent with previous observations (Greenblatt et al., 2018), the resulting chimeras showed a near-complete donor reconstitution as judged by CD45.2 staining, and normal frequencies of all major cell types (data not shown). However, Flt3L-supplemented cultures of primary BM cells yielded significantly reduced fractions of cDC2s (∼1.5-fold) and pDCs (∼2-fold) (Fig. 5C). Accordingly, the spleens of *Carm1*^Δ^ recipients manifested significant >2-fold reductions of cDC2s, pDCs and pDC-like CD8α^+^ Xcr1^-^ DCs (corresponding to transitional DCs or tDCs) (Bar-On et al., 2010; Leylek et al., 2019) (Fig. 5D,E). Notably, cDC1s were not reduced, suggesting a subset-specific role of *Carm1* in DC differentiation. Consistent with pDC reduction, the production of IFN-I in response to TLR9 ligand CpG oligonucleotides was profoundly reduced in *Carm1*^Δ^ recipients, whereas IFN-I response to the TLR3/MDA-5 ligand poly-I:C was normal (Fig. 5F).

To ensure that the observed phenotypes were cell-intrinsic, we established competitive chimeras using *Carm1*^Δ^ or control *Carm1*^fl/fl^ donor BM mixed with CD45.1^+^ wild-type competitor BM. The reconstitution by *Carm1*^Δ^ BM was ∼2-fold lower than by control BM, so that the ratio of *Carm1*^Δ^:control donor-derived cells was ∼0.5 in the total BM and in Flt3L cultures derived from it (Fig. S5C,D). However, the fractions of donor-derived cDC2s and pDCs, but not of cDC1s, were significantly lower in these cultures (Fig. S5C). Among primary cells, a strong reduction of *Carm1*^Δ^ donor-derived T cells was observed in these competitive settings, likely reflecting aberrant thymocyte development (Kim et al., 2004); importantly, however, a significant reduction was also observed in pDCs in the BM (Fig. S5E) and in primary splenic cDC2s, pDCs and tDCs but not cDC1s (Fig. S5F). To further explore the role of *Carm1* in DC development, we performed bulk RNA-Seq on CD45.2^+^ donor-derived DC subsets isolated from spleens of the recipients of *Carm1*^Δ^ or *Carm1*^fl/fl^ BM. Principal component analysis (data not shown) and pairwise comparisons (Fig. S5G) showed relatively few differences between Carm1-deficient and control DCs, with the most prominent differences observed in pDCs. Furthermore, both cDC2s and pDCs showed an aberrant upregulation of the cDC1 subset-specific gene signature (Fig. S5H). Collectively, these data suggest that *Carm1* is specifically required for optimal differentiation of cDC2s and especially pDCs, providing an *in vivo* validation for this candidate from the screen.

### Negative regulators of mTOR signaling are essential for DC differentiation

The mTOR signaling pathway, and specifically its major signaling complex mTORC1, coordinates cellular responses to nutrients and growth signals including the PI3K/Akt pathway activated by Flt3 (Kim and Guan, 2019; Liu and Sabatini, 2020). Its activation is dampened by constitutive negative regulators including the TSC and GATOR1 complexes, as well as by context-dependent negative regulators such as the FLCN-FNIP complex (Fig. 6A). Among the genes with highest CRISPTimeR scores in progenitors were *Tsc1* and *Tsc2*, encoding the two subunits of the TSC complex; as well as *Nprl2*, *Nprl3* and *Depdc5*, encoding the three subunits of the GATOR1 complex (Fig. 6B). Accordingly, the downstream negative effector of the PI3K/mTOR signaling, *Foxo3*, also had a high score whereas genes encoding AKT1 and mTOR itself had low scores in progenitors (Fig. 6B). This result is consistent with the expected positive role of mTOR in Flt3L-driven proliferation of progenitors. Strikingly, the opposite was observed in DCs, in which TSC and GATOR1 components were among the lowest-scoring genes. *Foxo3* also had a relatively low score, as did the genes encoding two other negative regulators of mTOR signaling, *Strada* and *Flcn* (encoding folliculin). This was also observed in pDCs and cDC2s, with *Tsc1* and *Tsc2* showing the lowest scores in all DC samples (Fig. 6C). The observed switch from high scores in progenitors to low scores in DCs was in contrast to *Pten*, which had high score in both samples (Fig. 3F).

**Figure 6.**
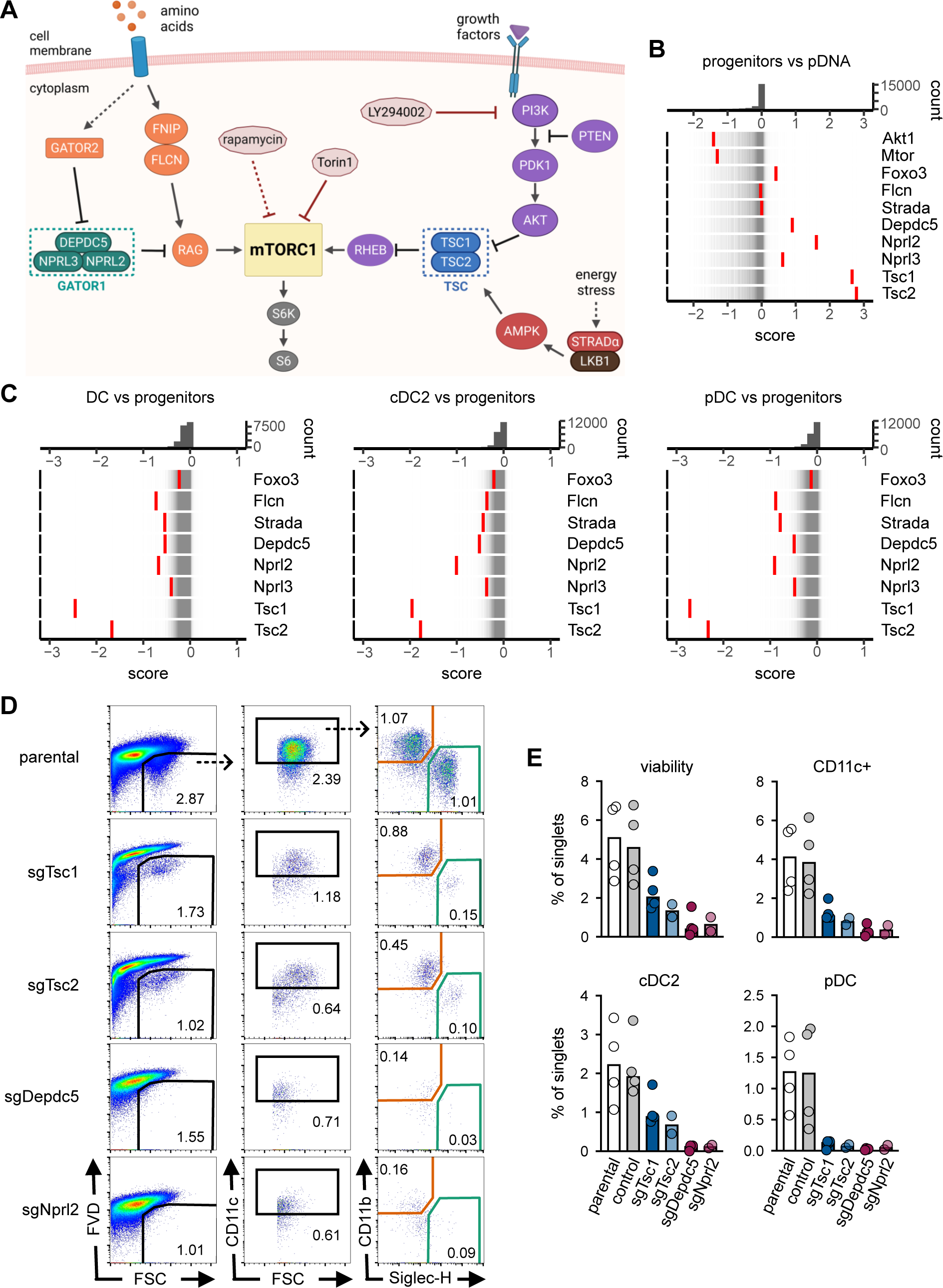
TSC and GATOR1 complexes inhibit DC differentiation. (A) Schematic of the mTOR signaling pathway and its regulators, including constitutive negative regulators (TSC and GATOR1 complexes), context-dependent activators or inhibitors (the FNIP/FLCN complex and STRADA), and small molecule inhibitors. (B-C) Rug plots of gene distribution between indicated pairs of library samples, as measured by the CRISPTimeR score. Top panels show the distribution of scores; bottom panels show individual genes (red) against the distribution of all genes (grey). (B) Rug plots of select components and inhibitors of the mTOR signaling pathway in Hoxb8-FL progenitors vs the sgRNA library (pDNA). (C) Rug plots of select inhibitors of the mTOR signaling pathway in Hoxb8-FL-derived DCs vs progenitors. (D-E) Differentiation capacity of Hoxb8-FL cells with targeted components of TSC and GATOR1 complexes. Parental Hoxb8-FL cells or cells transduced with control sgRNA or sgRNAs targeting indicated genes encoding TSC or GATOR1 subunits were differentiated into DCs and analyzed by flow cytometry as in Fig. S3B. (D) Sequential gating of single cells highlighting viable cells, CD11c^+^ DCs, and SiglecH^+^ pDCs and CD11b^+^ cDC2s. Numbers indicate the fraction of cells in the gate among single cells. Representative of two sgRNAs per gene. (E) Frequencies of DC subsets among single cells. Symbols represent separate differentiation experiments (n = 3); bars represent the median.

The requirement for TSC and GATOR1 was confirmed by transduction of Hoxb8- FL cells with individual sgRNAs targeting their subunits, which did not affect progenitor growth (Fig. S6A) but essentially abolished the differentiation of all DCs (Fig. 6D,E). Time course analysis of differentiating TSC- or GATOR1-deficient Hoxb8-FL cells showed that DC progenitors underwent massive apoptosis starting on day 4 (Fig. S6B), coinciding with the onset of terminal DC differentiation. Accordingly, DC progenitors were detectable on days 3-4 but failed to produce mature DC subsets by days 5-7 (Fig. S6C). Thus, TSC and GATOR1 complexes may restrict progenitor growth but are specifically required for DC differentiation, particularly for the survival of DC during terminal differentiation.

### Reduced mTOR signaling during DC differentiation

The observed role of mTOR inhibitory complexes prompted us to test the role of mTOR signaling in DC differentiation vs progenitor growth. First, we tested the effect of Torin1 (a direct inhibitor of the mTOR kinase) and rapamycin (an indirect inhibitor of the mTORC1 complex) on the growth and differentiation of Hoxb8-FL cells. As expected, both drugs inhibited the growth of undifferentiated Hoxb8-FL progenitors within 3-4 days in the nanomolar range (Fig. 7A,B). In contrast, they did not inhibit and even increased the differentiation of Hoxb8-FL cells into pDCs or cDCs in the same range (Fig. 7C). Notably, LY294002 (an inhibitor of PI3K) inhibited progenitor proliferation in the low micromolar range, with a near-complete inhibition at 10 μM (Fig. 7A,B). The same concentrations of LY294002 also enhanced DC differentiation whereas higher concentrations completely inhibited it, likely due to the block of PI3K signaling through other pathways. Treatment with Torin1 did not rescue the differentiation of TSC- or GATOR1-deficient cells, but improved their differentiation in a dose-dependent manner (Fig. S7A). Next, we confirmed a decrease in mTOR activity during the differentiation of Hoxb8-FL cells by Western blot for phosphorylated p70S6 kinase (p-p70S6K) a downstream effector of the mTORC1 complex (Fig. S7B,C), and by intracellular staining for phosphorylated S6 (p-S6) (Fig. 7D) the primary substrate of p70S6K (Fig. 6A). The levels of p-S6 were decreased in nearly all cDC2s and the majority of pDCs, although a fraction of pDCs retained high p-S6 levels (Fig. 7D); the reason for this heterogeneity remains to be investigated. The staining also confirmed higher p-S6 levels and hence increased mTOR activity in TSC- and GATOR1- deficient progenitors (Fig. S7D). Thus, DC differentiation is accompanied by a decrease in mTOR signaling, which is required for the process and is mediated by non-redundant activities of TSC and GATOR1 complexes.

**Figure 7.**
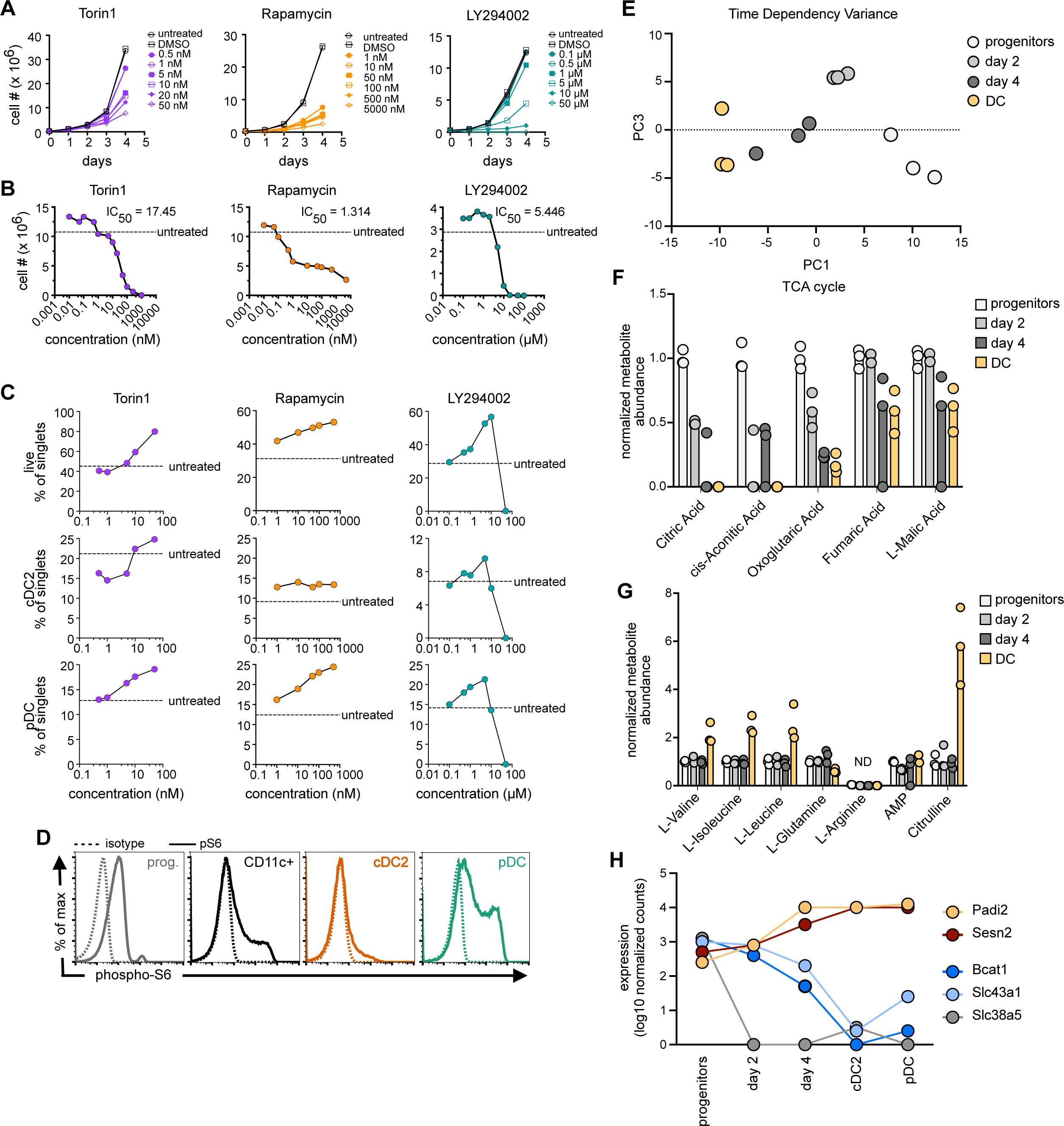
mTOR signaling is switched off during DC differentiation. (A-B) The growth of undifferentiated Hoxb8-FL cells in the presence of mTOR inhibitors Torin1 or rapamycin, or the PI3K inhibitor LY294002. Representative graphs of at least two experiments are shown. (A) Growth curves in the presence of indicated inhibitor concentrations. (B) Cell numbers on day 4 of growth in the presence of indicated inhibitor concentrations; the 50% inhibitory concentration (IC50) and the number in an untreated control culture (dashed line) are indicated. (C) The differentiation of Hoxb8-FL cells in the presence of mTOR inhibitors. Cells were differentiated for 7 days in the presence of the increasing inhibitor concentrations and stained by flow cytometry. The frequencies of viable cells, cDC2s and pDCs among single cells were determined. Representative of 2-4 experiments. (D) Intracellular staining for phospho-S6 in live undifferentiated Hoxb8-FL progenitors (prog.) or differentiated CD11c^+^ DCs, cDC2s or pDCs. Dashed histograms indicate isotype controls. Representative staining profiles of two experiments are shown. (E-G) Metabolomic profile of undifferentiated (progenitor), differentiating (day 2 and day 4) and bulk differentiated Hoxb8-FL cells (DC). Symbols represent biological replicates (n = 3); bars represent the median. (E) Principal component analysis of global metabolite profile of Hoxb8-FL cells at different stages of differentiation. (F) Abundance of metabolites of the tricarboxylic acid (TCA) cycle in Hoxb8-FL cells at different stages of differentiation. (G) Abundance of branched-chain amino acids (BCAAs) and of select metabolites of the urea cycle in in Hoxb8-FL cells at different stages of differentiation. (H) Expression of select molecules involved in mTOR activation by amino acids during HoxB8-FL cell differentiation as determined by RNA-Seq (Fig. 2). Shown are average log10 expression levels of indicated genes on days 0-4 of differentiation and in mature cDC2s and pDCs on day 7.

To correlate the observed switch of mTOR signaling with metabolic activity during DC differentiation, we performed metabolic profiling of differentiating HoxB8-FL cells on days 0 (progenitors), 2, 4 and 7 (DCs). Of 147 metabolites measured by mass spectrometry, 117 could be detected in more than one sample. Principal component analysis (PCA) of the resulting metabolite levels showed the expected alignment of samples along their differentiation trajectory (Fig. 7E). We then identified metabolites that were significantly changed in differentiated DCs compared to progenitors (Fig. S7E). Most of these metabolites were enriched in progenitors and corresponded to multiple pathways including pyrimidine metabolism and the TCA cycle, a major biosynthetic hub activated by mTOR (Fig. S7F). Indeed, TCA cycle intermediates were progressively reduced during DC differentiation (Fig. 7F), consistent with the reduction of mTOR signaling.

The majority of metabolites upregulated in DCs over progenitors comprised amino acids (Fig. S7E). This was unexpected, given that amino acid availability is a key factor activating the mTOR pathway. We therefore focused on amino acids that serve as major activators as well as targets of mTOR signaling, namely branched chain amino acids (BCAA: valine, isoleucine and leucine), glutamine and arginine (Liu and Sabatini, 2020; Mossmann et al., 2018). BCAA were increased ∼2-fold in differentiated DCs (Fig. 7G); however, genes encoding BCAA transporter SLC43A1 and BCAA-ketoacids interconverting enzyme BCAT1, both of which facilitate BCAA-mediated mTOR activation, were dramatically downregulated (Fig. 7H). Conversely, the gene encoding SESN2, an inhibitor of leucin-induced mTOR signaling, was strongly induced during DC differentiation (Fig. 7H). Glutamine was reduced ∼2-fold in DCs (Fig. 7G), and accordingly, the transcript encoding glutamine transporter SLC38A5 was rapidly downregulated during differentiation (Fig. 7H). While arginine was undetectable in our analysis, we noted a strong induction in DCs of citrulline, a product of arginine conversion in the urea cycle as well as by peptidyl arginine deiminase (PADI) enzymes (Fig. 7G). The only gene for a PADI enzyme expressed in HoxB8-FL cells by RNA-Seq was *Padi2*, and its expression was strongly upregulated during DC differentiation (Fig. 7H). Overall, metabolomic profiling of DC differentiation is consistent with a progressive reduction of mTOR signaling, and suggests that the availability of key amino acids may contribute to it.

## Discussion

We applied the conditionally immortalized, cytokine-dependent progenitor cell line Hoxb8-FL to interrogate the process of DC development at genome-wide level. This model allows the generation of two major DC subsets in a process driven by the biologically relevant growth factor Flt3L. We confirmed that the resulting pDCs and cDC2s are closely related to their primary counterparts with respect to their transcriptome and chromatin profile. Of note, Hoxb8-FL-derived DCs have not received the Notch2- mediated signal that guides terminal cDC2 differentiation (Lewis et al., 2011) and facilitates the emergence of cDC1s (Kirkling et al., 2018). As such, these cells apparently represent relatively immature cDCs, particularly in their open chromatin profile. We also showed that DC differentiation from Hoxb8-FL closely resembles the differentiation of DCs *in vivo* as characterized by inducible lineage tracing (Feng et al., 2022). In particular, the uniform expression of Cx3cr1 and Csf1R on differentiating progenitors is similar to the phenotypes of early DC progenitors (Fogg et al., 2006; Naik et al., 2007; Onai et al., 2007) and supports the derivation of both pDCs and cDCs from these cells. This is also consistent with a cell-intrinsic mechanism of cDC/pDC lineage specification based on the self-amplifying activity of TCF4 and its inhibitor ID2 (Grajkowska et al., 2017). These data validate Hoxb8-FL cells as a useful *in vitro* model of DC development, and support the derivation of pDCs and cDCs from common progenitors *in vivo*, even in the absence of terminal cDC1 differentiation.

We performed a genome-wide CRISPR/Cas9 dropout screen in Hoxb8-FL cells to identify regulators of progenitor proliferation and DC differentiation. The sgRNA library comprised only four sgRNAs per gene; thus, a suboptimal design of only 1-2 sgRNAs per gene would result in a false negative. Indeed, many known regulators of DC differentiation such as *Irf8* and *Tcf4* were missed in the screen; in the case of *Tcf4*, all sgRNAs in the library targeted the most 5’ exon, thus sparing the equally functional transcript from the downstream start site (Grajkowska et al., 2017). On the other hand, positive hits were identified with high confidence, as supported by i) hits on many well-known positive (e.g. *Flt3*, *Spi1*) and negative (e.g. *Pten*, *Bcor*) regulators; ii) hits on multiple subunits of the same protein complexes such as TSC, GATOR1, NPC and GPI transamidase; iii) validation of two newly identified regulators (*Npc1* and *Carm1*) by conventional gene targeting *in vivo*. We also explored the DC subset specificity of identified candidates by interrogating them separately in pDCs and cDC2s, although this analysis was limited by low numbers of sorted cells. The vast majority of candidates behaved similarly in both subsets, suggesting that they represent general regulators of DC differentiation. Nevertheless, sgRNAs for several known regulators of DC differentiation (such as *Irf2* and *Bcl11a*) showed a more pronounced depletion in the respective subset, supporting the specificity of the screen.

The results yielded multiple genes that were dispensable for progenitor proliferation but facilitated DC differentiation. Some of these genes appear to control differentiation of multiple immune and/or non-immune cell types; thus, lysine methyltransferase *Setdb1* is broadly required for hematopoietic differentiation (Koide et al., 2016) and T cell effector differentiation (Adoue et al., 2019), whereas the GPI transamidase complex genes control embryonic stem cells (ESC) differentiation (Li et al., 2018). The role of other identified genes in cell differentiation has not been previously reported, and should provide a useful resource for future studies in the field. In addition, we identified and confirmed *in vivo* a hitherto unknown role of the NPC cholesterol transporter complex in DC differentiation. NPC1 was not absolutely required for DC differentiation but facilitated it in vivo. The results help explain the reported reduction of T cell priming in *Npc1*-deficient mice, even though the antigen presentation capacity of DCs was intact (Bosch et al., 2013). The mechanism whereby the NPC complex facilitates DC differentiation remains to be elucidated, and may include its recently proposed role as a regulator of STING-mediated interferon signaling (Chu et al., 2021).

The screen also identified protein methyltransferase *Carm1* as a candidate positive regulator of DC differentiation. This was unexpected given that *Carm1* is expressed ubiquitously and was shown to be dispensable for adult hematopoiesis, although its function in DCs was not studied (Greenblatt et al., 2018). Indeed, a re-evaluation of *Carm1*-deficient hematopoietic cells revealed a significant, cell-intrinsic reduction of pDCs and cDC2s, the two DC subsets interrogated in the screen. In contrast, myeloid cells in general and cDC1s in particular were unaffected, revealing a DC subset-specific activity of *Carm1*. CARM1 is a pleiotropic enzyme that methylates multiple substrates and can affect differentiation through co-activation of transcription and other mechanisms. Interestingly, CARM1 was shown to methylate MED12 in the Mediator transcriptional complex, and this activity requires JMJD6, another hit from the screen (Gao et al., 2018). Moreover, the CARM1-JMJD6 complex on the Mediator target genes includes BRD4, an essential regulator of pDC differentiation (Grajkowska et al., 2017). Another relevant substrate of CARM1 is p300 (Chen et al., 2000), through which CARM1 controls multiple transcriptional pathways such as NF-κB (Covic et al., 2005). In any case, our results identify CARM1 as an important mediator of DC differentiation and provide a critical validation of the screen *in vivo*.

A notable feature of the Hoxb8-FL cell system is the sharp separation between the proliferation and the differentiation stages, which facilitates independent analysis of the two processes (Grajkowska et al., 2017). Indeed, sgRNAs for many hits were enriched in progenitors and subsequently depleted in DCs, suggesting that these molecules restrict proliferation and promote differentiation. The most striking examples of this “switch” behavior were provided by subunits of the TSC and GATOR1 complexes, two well-established inhibitors of mTOR signaling pathway. These results, together with additional dissection of the mTOR activity in Hoxb8-FL cells, suggested that TSC- and GATOR1- mediated inhibition of mTOR activity occurs at the onset of DC differentiation and is required for the survival of differentiating DCs. This was in contrast to the more upstream activity of the PI3K pathway, which promoted both progenitor proliferation and DC differentiation, and whose inhibitor *Pten* opposed both processes. The specific non-redundant activity of TSC and GATOR1 in promoting cell differentiation is not common to all cell types, as exemplified by findings in other genome-wide screens: thus, GATOR1 promoted both the proliferation and the differentiation of ESCs, whereas TSC inhibited both processes (Li et al., 2018). On the other hand, mTOR is required for optimal effector differentiation of key immune cell subsets such as regulatory T cells and memory T cells (Powell and Delgoffe, 2010).

Our data help reconcile conflicting observations on the role of PI3K/mTOR signaling in DC differentiation, such as the apparently opposite roles of *Pten* and *Tsc1* in the process (Sathaliyawala et al., 2010; Wang et al., 2013). They also explain why rapamycin inhibits in vitro DC development from hematopoietic progenitors (Sathaliyawala et al., 2010; van de Laar et al., 2010) but can be applied to promote the development of monocyte-derived DCs (Erra Diaz et al., 2020; Macedo et al., 2012), because the former but not the latter cultures involve extensive progenitor proliferation. More importantly, they identify the TSC/GATOR1-dependent downregulation of mTOR signaling as a key molecular switch between progenitor expansion and terminal DC differentiation. In DCs, unlike in other lineages such as lymphocytes or granulocytes, both processes are driven by the same cytokine receptor, Flt3. Our results suggest that a selective shutdown of the mTOR pathway redirects Flt3 signaling to induce DC differentiation, revealing a selective “interpretation” of the same growth factor signal by differentiating cells. In addition to ensuring DC survival, the reduction of mTOR signaling in mature DCs is likely important for their eventual activation by microbial stimuli, which induce mTOR signaling leading to cytokine production and antigen presentation (Cao et al., 2008; Fekete et al., 2020). The results also raise an important question of the molecular mechanism that induces TSC/GATOR1-dependent mTOR inhibition downstream of Flt3. One candidate for this role is FLCN, which was identified in our screen as a positive regulator of differentiation, but appeared neutral for proliferation. Unlike constitutive inhibitors of mTOR such as TSC and GATOR1, FLCN is a context-dependent mTOR inhibitor (Ramírez-Reyes et al., 2021) and thus may control the switch from mTOR activation to inhibition. Interestingly, NPC1 was also identified as a negative regulator of mTOR signaling in some contexts (Castellano et al., 2017), suggesting that the NPC complex may facilitate DC differentiation in part by inhibiting mTOR. Our data also suggest that the availability of key amino acids, brought about by multiple genes, may contribute to the observed reduction of mTOR signaling. Whereas the role of these and other mechanisms remains to be identified in future studies, our results establish the switch in mTOR activity as a fundamental mechanism of Flt3-driven DC development.

## Acknowledgments

Supported by the NIH grants AI072571 and AI154864 (B.R) and GM126421 (M.T.B.), the Dr. Bernard Levine postdoctoral fellowship program in immunology (I.T.), the Damon Runyon postdoctoral fellowship DRG 2408-20 (N.M.A.), and by the Center for Experimental Immuno-oncology at Memorial Sloan Kettering Cancer Center and Comedy vs Cancer (C.M.L). We acknowledge the use of resources provided by NYU Metabolomics Laboratory, NYU Genome Technology Center (GTC), NYU Applied Bioinformatics Facility Laboratories (ABL) and the NYU High Performance Computing Facility (HPCF). GTC and ABL are Shared Resources partially supported by NIH grant P30CA016087 at the Laura and Isaac Perlmutter Cancer Center.

## Author contributions

I.T., P.-F.H., N.M.A., C.S., F.L. and T.C.R. performed experiments and analyzed results. L.S.L.-Z., C.M.L., E.E., A.K.-J., D.J., A.T. and U.O. developed analytical methods and analyzed results. M.T.B. and S.D.N. provided essential reagents. I.T. and B.R. conceived the project, analyzed results and wrote the manuscript with input from all authors.

## Methods

### Mice

All animal studies were performed according to the investigator’s protocol approved by the Institutional Animal Care and Use Committee of New York University School of Medicine. Mice with the deletion of *Carm1* in hematopoietic cells have been generated by crossing the LoxP-flanked conditional allele of *Carm1* (*Carm1*^flox^) with pan-hematopoietic *Vav1*-Cre transgenic deleter strain (Greenblatt et al., 2018). The resulting *Carm1*^fl/fl^ *Vav1*-Cre or littermate control *Carm1*^fl/fl^ mice were used as BM donors for hematopoietic reconstitution. Total BM (2×10^6^ cells) were injected i.v. into lethally irradiated male 8-week CD45.1 congenic C57BL/6 mice (B6.SJL-*Ptprc^a^Pepc^b^*/BoyCrCrl, Charles River). To establish competitive chimeras, donor BM was mixed 1:1 with the BM from CD45.1 congenic mice, and 2×10^6^ cells were used for reconstitution as above. *Npc1*^- /-^ mice (B6.C-*Npc1^m1N^*/GarvJ; Strain #030097) were purchased from the Jackson Laboratory, crossed to wild-type C57BL/6 mice and intercrossed to generate the *Npc1*^-/-^ and *Npc1*^+/+^ littermate controls used in this study. Male and female mice were used at around 6-7 wk of age, prior to the appearance of neurological symptoms. Hematopoietic reconstitution with the BM from *Npc1*^-/-^ or control *Npc1*^+/+^ mice was performed as above.

### Cell culture and inhibitor treatments

The FLT3L-secreting clone of the C57BL/6- derived B16 melanoma cell line (Mach et al., 2000) was used to produce FLT3L- containing supernatants and was cultured in DMEM supplemented with 10% FCS, 1% L-glutamine, 1% sodium pyruvate, 1% MEM-NEAA, and 1% penicillin/streptomycin. The murine progenitor Hoxb8-FL cell line was derived and cultured as described (Redecke et al., 2013). In brief, the cells were grown in RPMI supplemented with 10% FCS, 10% supernatant from the Flt3L-producing B16 cell line, and 10 μM β-estradiol. The cells were induced to differentiate into DCs by washing and replating in RPMI supplemented with 10% charcoal-stripped FCS and 10% Flt3L supernatant. The resulting differentiated cells were collected by scraping on day 7. Day 2 and day 4 samples were similarly collected two and four days post differentiation induction. For chemical inhibition experiments, cells were cultured with indicated concentrations of TP-064 (Tocris Bioscience), EZM2302 (DC Chemicals), JQ1 (Sigma-Aldrich), Torin1 (Calbiochem), rapamycin (Calbiochem) or LY294002 (Cell Signaling Technology), all dissolved in DMSO. All inhibitors were added with fresh medium once at the start of treatment. The highest matching concentration of DMSO or medium alone were used as controls.

### Primary cell preparations

Spleens were minced and digested with collagenase D (1 mg/ml) and DNase I (20 µg/ml) in full DMEM for 30–60 min at 37°C. Tissues were pressed through nylon 70 µm cell strainer to yield single-cell suspensions and then subjected to red blood cell (RBC) lysis (155 mM NH4Cl, 10 mM NaHCO3, and 0.1 mM EDTA) for 5 min at room temperature before being filtered. BM was prepared by flushing femurs and tibias with PBS using a 27-gauge needle before RBC lysis. Lymph nodes were pressed through a 70-µm cell strainer to generate single-cell suspensions. Peripheral blood (PB) was obtained from submandibular vein and subjected to RBC lysis for 5 minutes, followed by 3 min, at room temperature (RT).

### FLT3L-driven DC differentiation of primary BM cultures

Single cell suspensions of primary BM cells were obtained as described above. Cells were plated in 24-well plates at a density of 2×10^6^ cells in 2 mL per well, cultured in full DMEM (10% FCS, 1% L-glutamine, 1% sodium pyruvate, 1% MEM-NEAA and 1% penicillin/streptomycin, 55 μM 2-mercaptoethanol) supplemented with 10% supernatant from the cultured B16-FLT3L cell line, and harvested 7 days later.

### Generation of Hoxb8-Cas9 cell line

To express Cas9 constitutively in Hoxb8-FL cells, we used lentiCas9-Blast vector that confers blasticidin resistance (Sanjana et al., 2014) (Addgene #52962). VSV-G pseudotyped lentiviral particles were produced using transient transfection in 293T cells. Undifferentiated Hoxb8-FL cells were spinoculated with the Cas9 vector and batch selected with blasticidin. The resulting population was cloned by limiting dilution and screened for the expression of Cas9 by Western blotting with anti- Cas9 antibody (Cell Signaling Technology). A clone with the highest expression of Cas9 and intact differentiation capacity was chosen for gene targeting.

### Targeting of candidate genes in Hoxb8-FL cells

Selected candidate genes identified in the screen were independently targeted using the CRISPR/Cas9 approach. SgRNAs targeting each gene (derived from the Brie library) were cloned into the lentiGuide-Puro lentiviral vector (Sanjana et al., 2014) (Addgene #52963) that confers puromycin resistance. sgRNAs used for targeting are shown in the Supplemental Table. VSV-G pseudotyped lentiviral particles were produced using transient transfection in 293T cells. Undifferentiated Cas9-expressing Hoxb8-FL cells were spinoculated with each of the sgRNA-containing vectors and batch selected with puromycin for subsequent analysis.

### Brie library lentivirus production and titration

The mouse CRISPR knockout pooled library Brie (Doench et al., 2016) was purchased from Addgene (#73633) and amplified according to the depositor’s Library Amplification Protocol. The amplified library was sequenced according to depositor’s Sequencing Protocol to confirm the maintenance of representation. VSV-G pseudotyped lentiviral particles were produced using transient transfection in 293T cells and the lentiviral stock was concentrated using the Lenti-X^TM^ Concentrator (Takara Bio) according to the manufacturer’s recommendations. Optimal transduction conditions for the screening were determined in order to achieve ∼30% transduction efficiency, corresponding to a multiplicity of infection (MOI) of ∼0.5. Briefly, undifferentiated Hoxb8-FL-Cas9 cells were plated in 6-well plates (5x10^5^ cells per well) and spinoculated with increasing volumes of the concentrated Brie lentivirus. The next day the cells were split 1:2 and replated, and 5 μg/mL puromycin was added to one well of each condition 24 h later. Wells with uninfected cells with or without puromycin served as controls. Cells were counted two days post selection to determine transduction efficiency by comparing survival with and without puromycin selection. The volume of virus that yielded ∼ 30% transduction efficiency was used for the screening.

### Brie library CRISPR screen

In order to achieve a representation of ∼400 cells per sgRNA in the library post-selection, 1.1x10^8^ undifferentiated Hoxb8-FL-Cas9 cells were distributed to 6-well plates with 5x10^5^ cells per well. Cells were spinoculated with the pre- determined amount of virus and 5 μg/mL puromycin was added 48 h later. Transduced cells were maintained in puromycin throughout the screen and were split every two days to prevent overconfluence. Samples of 5x10^7^ cells were obtained 5 days post selection (progenitors_0) and 11 days post selection on the day of differentiation induction (progenitors). A bulk sample of 10^8^ differentiated cells (DC) was obtained at the end of differentiation and prior to sorting. Hoxb8-FL-derived DCs were sorted as follows: CD11c^+^ CD11b^+^ B220^-^ for cDC2s (4.6x10^6^) and CD11c^+^ CD11b^-^ B220^+^ for pDCs (3x10^6^). All sample cell pellets were stored at -80°C until genomic DNA (gDNA) isolation.

gDNA was extracted using the QIAamp Blood Maxi/Midi kit (QIAGEN) according to the manufacturer’s instructions and concentration was measured by Nanodrop (ThermoFisher). Libraries were prepared and purified with AMPure XP-PCR Purification kit (Beckman) according to the depositor’s Sequencing Protocol (Addgene #73633) For “progenitors_0” and “progenitors” samples, 170-190 μg DNA corresponding to ∼400x coverage of the library were amplified in 17-19 parallel PCR reactions using 10 μg per reaction. For the “DC” sample, 38 PCR reactions were similarly setup for the amplification of 380 μg of DNA corresponding to ∼ 800x coverage of the library. For sorted cDC2 and pDC samples library coverage was limited due to low cell numbers. One PCR reaction was setup for each sorted subset for the amplification of ∼7-9 μg genomic DNA.

Paired-end sequencing for 50 cycles was performed using Illumina HiSeq4000 sequencer. For the analysis of demultiplexed sequencing files python count_spacers.py was adjusted as needed and run to generate sgRNA read counts and relevant statistics for each sample (Joung et al., 2017). Read counts from all samples were joined in a data frame and normalized by library size using the following formula:

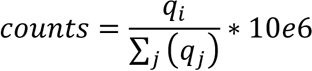

Where *q*_*i*_ corresponds to the reads mapped to a specific sgRNA sequence, and ∑(*q_j_*) corresponds to the total number of mapped reads to sgRNAs.

### Identification of significantly enriched/depleted sgRNAs by empirical false discovery rate (FDR)

After normalizing for library size, the read counts of each sample were compared with the counts of the pDNA sample, producing a log2 fold change table. To determine significance of the fold change of each sgRNA, a null distribution was built for each sample using the 1000 non-targeting sgRNAs included in the library. Single tailed critical values were obtained for an α of 0.05 for both tails of the distribution and classified each sgRNA as enriched, depleted or non-changing based on the rejection areas generated.

### Identification of significantly enriched/depleted genes by mixed linear model

After normalization, a mixed linear model was used to evaluate gene enrichment or depletion as described (Garipler et al., 2022). Briefly, the model has the following formula:

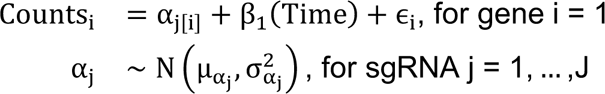

Where Counts_i_ is the log counts of gene i ; α_<[:]_ is the intercept for sgRNA j of gene i ; β_A_ is the overall estimate for Time and ɛ is the error. A random intercept model was chosen to account for the variability in the sgRNAs initial observed counts. Hypothesis testing was done using likelihood ratio test via ANOVA. The model was compared to a reduced version that does not account for time. P-value adjustment was performed with the Benjamini-Hochberg Procedure. CRISPTimer Score was then calculated by multiplying the estimate by the -log10 of the adjusted p value. This analysis was performed for each sample.

### Brie library CRISPR screen data downstream analysis

Mixed linear models were fitted using the lmer function of lme4. Dataframe import, tidying and manipulation were done using the readr, tidyr and dplyr packages from the tidyverse (1.3.1). Violin plots, rug plots, dot plots and histograms were made using ggplot2 R package. Shannon diversity index was calculated using vegan (2.4-2) R package, and plot was made using R base plot. Enrichment analysis in Fig.S3F was performed using Enrichr (Kuleshov et al., 2016) against the Gene Ontology Biological Processes version 2021 gene-set library and bar charts were generated using the enrichment analysis visualizer Enrichr Appyter.

### Flow cytometry

Single-cell suspensions of cultured DCs (Flt3L-supplemented BM cultures) or primary cells (splenocytes, BM or blood cells) were prepared as described above and stained for multicolor analysis with the indicated fluorochrome- or biotin-conjugated antibodies. For progenitor analysis, BM cells were stained with a cocktail of antibodies to lineage markers (B220, CD11b, TCRb, Gr1, NK1.1, and Ter119). With the exception of Hoxb8-FL cells, staining of surface molecules with fluorescently labeled antibodies was performed for 20 min at 4°C in the dark. Hoxb8-FL cells were stained for 15 min at room temperature. Intracellular staining of cells with phospho-S6 (eBioscience) was performed for 20 min at room temperature using the Intracellular Fixation & Permeabilization Buffer Set (eBioscience) according to the manufacturer’s recommendations. For apoptosis assessment, Hoxb8-FL cells were stained using the FITC Annexin V Apoptosis Detection Kit with 7-AAD (BioLegend) according to the manufacturer’s instructions. Samples were acquired on Attune NxT (Invitrogen) using Attune NxT software and further analyzed with FlowJo software.

### Western blot

Cell pellets of Hoxb8-FL-derived DCs were lysed on ice for 30 min in RIPA lysis buffer (50 mM Tris-HCl pH 8.0, 150 mM NaCl, 1% Triton X-100, 0.1% SDS, and 0.5% sodium deoxycholate) supplemented with Halt™ Protease Inhibitor Cocktail (Thermo Scientific) and boiled for 10 min in the presence of 4x SDS sample loading buffer. For mTOR signaling pathway-related antibodies, cell pellets of Hoxb8-FL cells at different stages of differentiation were directly lysed in SDS sample loading buffer and boiled for 10 min. Lysates from equal cell numbers were analyzed by SDS-PAGE followed by western blotting with total protein (for phosphorylated proteins) or β-actin used as loading control.

### In vivo IFN response

To induce an IFN response in vivo, *Carm1*^Δ^ (*Carm1*^fl/fl^ *Vav1*-Cre) or *Carm1*^fl/fl^ control mice were injected i.v. with 5 µg of CpG-A (ODN 2216; InvivoGen) complexed with DOTAP (30 µl DOTAP/100 µl total volume; Roche), and blood was collected 6 h later. Three weeks later the same mice were injected i.p. with 250 µg of poly(I:C) (HMW; InvivoGen) and blood was collected 6 h later. IFN-α concentration in the sera was determined by ELISA (VeriKIne HS mouse IFN-α all subtype; PBL Assay Science) according to the manufacturer’s recommendations.

### Cell sorting and sample preparation for RNA-Seq and ATAC-Seq

For transcriptome profiling of *Carm1*-deficient DCs, chimeras reconstituted with the BM of *Carm1*^Δ^ or *Carm1*^fl/fl^ littermate controls (n= 3) were used as a source of cells. Splenic DCs were first enriched by positive selection using biotinylated anti-CD11c streptavidin microbeads and MACS columns (Miltenyi Biotec), and then sorted on FACSAria (BD) instrument. CD45.2^+^ DC populations were sorted as follows: cDC1 (CD11c^hi^ MHCII^+^ CD11b^-^ CD8α^+^ Xcr1^+^), cDC2 (CD11c^hi^ MHCII^+^ CD11b^+^ CD8α^-^) and pDC (SiglecH^+^ B220^+^). For RNA-seq, RNA from sorted cells was extracted using the RNeasy mini kit (QIAGEN) according to the manufacturer’s instructions.

For transcriptome and epigenetic profiling of Hoxb9-FL-derived DC development, Hoxb8- FL cells at different stages of differentiation were collected and sorted as follows: total single live cells for undifferentiated progenitors (day 0), day 2 and day 4 cells; CD11c^+^ CD11b^+^ B220^-^ for cDC2s and CD11c^+^ CD11b^-^ B220^+^ for pDCs. For RNA-Seq, RNA from sorted cells (∼10^6^) was extracted using the RNeasy mini kit (QIAGEN) and the QIAshredder homogenizer (QIAGEN) according to the manufacturer’s instructions. cDNA libraries were prepared using the Low input Clontech SMART-Seq HT with Nxt HT and paired-end sequencing was performed on Illumina NovaSeq 6000.

For ATAC-seq, sorted cells (1x10^5^) were washed with PBS and prepared as described (Buenrostro et al., 2015). Paired-end sequencing was performed in Illumina HiSeq 4000. All sequencing was performed at the Genome Technology Core at New York University School of Medicine.

### RNA-Seq data analysis

Sequencing reads were mapped to the reference genome (mm10) using the STAR aligner (v2.5.0c) (Dobin et al., 2013). Alignments were guided by a Gene Transfer Format (GTF) file. The mean read insert sizes and their standard deviations were calculated using Picard tools (v.1.126) (http://broadinstitute.github.io/picard). The read count tables were generated using HTSeq (v0.6.0) (Anders et al., 2015), normalized based on their library size factors using DEseq2 (Love et al., 2014), and differential expression analysis was performed. The Read Per Million (RPM) normalized BigWig files were generated using BEDTools (v2.17.0) (Quinlan and Hall, 2010) and bedGraphToBigWig tool (v4). To correct for global differences between primary and cultured cells, the removeBatchEffect function from limma (v.3.38.3) was used, by providing the cell source information (primary vs. Hoxb8- FL) as a factor. Gene set enrichment in Fig. 1E-F was performed first by using EdgeR (v.3.24.3) (Robinson et al., 2010) to calculate normalization factors and dispersion estimates. These calculations were used for rotation gene set testing (ROAST) (Wu et al., 2010) using the fry function to generate a two-sided directional p-value. log2FC values shown in barcode plots are those calculated using the EdgeR pipeline. Gene set enrichment analysis in Fig. S5H was performed using GSEA tool (Subramanian et al., 2005). Unique signature genes per conditions were found using the “findMarkers” function in iCellR R package (1.6.4) (Li et al., 2020) and the heatmap was generated using the heatmap.gg.plot function in iCellR. To compare the level of similarity among the samples and their replicates, we used two methods: principal-component analysis and Euclidean distance-based sample clustering. All the downstream statistical analyses and generating plots were performed in R environment (v3.1.1 or v3.5.3) (https://www.r-project.org/). Volcano and MA plots were made using ggplot2 R package and heatmaps using pheatmap or ComplexHeatmap (v1.99.7) R packages. Genomic track signals were generated using the Gviz R package (v1.18.2) using bigWigs described above.

### ATAC-Seq data analysis

All of the reads from the Sequencing experiment were mapped to the reference genome (mm10) using the Bowtie2 (v2.2.4) (Langmead and Salzberg, 2012) and duplicate reads were removed using Picard tools (v.1.126) (http://broadinstitute.github.io/picard/). Low quality mapped reads (MQ<20) were removed from the analysis. The read per million (RPM) normalized BigWig files were generated using BEDTools (v.2.17.0) (Quinlan and Hall, 2010) and the bedGraphToBigWig tool (v.4). Peak calling was performed using MACS (v1.4.2) (Zhang et al., 2008) and peak count tables were created using BEDTools. Differential peak analysis was performed using DESeq2 (Love et al., 2014). ChIPseeker (v1.8.0) (Yu et al., 2015) R package was used for peak annotations and motif discovery was performed using HOMER (v4.10) (Heinz et al., 2010). ngs.plot (v2.47) and ChIPseeker were used for TSS site visualizations and quality controls. To correct for global differences between primary and cultured cells, the removeBatchEffect function from limma (v.3.38.3) was used, by providing the cell source information (primary vs. Hoxb8-FL) as a factor. Gene set enrichment was performed first by using EdgeR (v.3.24.3) (Robinson et al., 2010) to calculate normalization factors and dispersion estimates. These calculations were used for rotation gene set testing (ROAST) (Wu et al., 2010) using the fry function to generate a two-sided directional p-value. log2FC values shown in barcode plots are those calculated using the EdgeR pipeline. Unique signature peaks per conditions were found using the “findMarkers” function in iCellR R package (1.6.4) (Li et al., 2020) and the heatmap was generated using the heatmap.gg.plot function in iCellR. To compare the level of similarity among the samples and their replicates, we used two methods: principal-component analysis and Euclidean distance-based sample clustering. The downstream statistical analyses and generating plots were performed in R environment (v3.1.1 or v3.5.3) (https://www.r-project.org/). Volcano plots were made using ggplot2 R package and heatmaps using pheatmap or ComplexHeatmap R packages. To generate peak-centered heatmaps, signal for each sample was quantified from bigWigs described above. Scores were generated using the computeMatrix function from deepTools2.0 (v3.21.1), with arguments “--referencePoint center --missingDataAsZero -b 1000 -a 1000 -bs 50”. For each condition, scores from each bin position were summarized across condition replicates. Genomic track signals were generated using the Gviz R package using bigWigs described above.

### Metabolite extraction and LC-MS/MS metabolomics

Hoxb8-FL cells at different stages of differentiation were collected (∼4 x 10^6^ cells per sample) as described above, washed in PBS and spun down. Metabolites were extracted from frozen cell pellets based on a previously described method (Pacold et al., 2016). Briefly, metabolite extraction was performed using an 80% methanol extraction buffer containing 500 nM isotopically labeled amino acid standard (MSK-A2, Cambridge Isotope Laboratories, Inc) alongside blank and standard controls at a ratio of 5 x 10^6^ cells per 1 mL extraction buffer. The cells were then homogenized with zirconium disruption beads (0.5 mm, RPI), centrifuged and a fixed volume (450 μL per sample) of the supernatant was speed vacuum concentrated to dryness at room temperature. The dried extracts were resolubilized in 50 μL LC-MS grade water. Samples were then subjected to liquid chromatography-mass spectrometry (LC-MS) analysis to detect and quantify known peaks. The LC column was a Millipore^TM^ ZIC-pHILIC (2.1 x150 mm, 5 μm) coupled to a Dionex Ultimate 3000^TM^ system and the column oven temperature was set to 25°C for the gradient elution. A flow rate of 100 μL/min was used with the following buffers; A) 10 mM ammonium carbonate in water, pH 9.0, and B) neat acetonitrile. The gradient profile was as follows; 80-20%B (0-30 min), 20-80%B (30-31 min), 80-80%B (31-42 min). Injection volume was set to 2 μL for all analyses (42 min total run time per injection).

MS analyses were carried out by coupling the LC system to a Thermo Q Exactive HF^TM^ mass spectrometer operating in heated electrospray ionization mode (HESI). Method duration was 30 min with a polarity switching data-dependent Top 5 method for both positive and negative modes. Spray voltage for both positive and negative modes was 3.5kV and capillary temperature was set to 320°C with a sheath gas rate of 35, aux gas of 10, and max spray current of 100 μA. The full MS scan for both polarities utilized 120,000 resolution with an AGC target of 3e6 and a maximum IT of 100 ms, and the scan range was from 67-1000 *m*/*z*. Tandem MS spectra for both positive and negative mode used a resolution of 15,000, AGC target of 1e5, maximum IT of 50 ms, isolation window of 0.4 m/z, isolation offset of 0.1 m/z, fixed first mass of 50 m/z, and 3-way multiplexed normalized collision energies (nCE) of 10, 35, 80. The minimum AGC target was 1e4 with an intensity threshold of 2e5. All data were acquired in profile mode.

### Hybrid metabolomics data processing and relative quantification of metabolites

The resulting Thermo^TM^ RAW files were converted to mzXML format using ReAdW.exe version 4.3.1 to enable peak detection and quantification. The centroided data were searched using an in-house python script Mighty skeleton version 0.0.2 and peak heights were extracted from the mzXML files based on a previously established library of metabolite retention times and accurate masses adapted from the Whitehead Institute (Chen et al., 2016), and verified with authentic standards and/or high resolution MS/MS spectral manually curated against the NIST14MS/MS (Simon-Manso et al., 2013) and METLIN (2017) (Smith et al., 2005) tandem mass spectral libraries. Metabolite peaks were extracted based on the theoretical *m*/*z* of the expected ion type e.g., [M+H]^+^, with a ±5 part-per-million (ppm) tolerance, and a ± 7.5 second peak apex retention time tolerance within an initial retention time search window of ± 0.5 min across the study samples. The resulting data matrix of metabolite intensities for all samples and blank controls was processed with an in-house statistical pipeline Metabolyze version 1.0 and final peak detection was calculated based on a signal to noise ratio (S/N) of 3X compared to blank controls, with a floor of 10,000 (arbitrary units). For samples where the peak intensity was lower than the blank threshold, metabolites were annotated as not detected, and the threshold value was imputed for any statistical comparisons to enable an estimate of the fold change as applicable. Due to the inherent size differences of mature DCs compared to progenitors, we *a priori* specified to sum-normalize the metabolic profile of each sample. This was accomplished by taking the sum of all detected metabolite signals in a given sample while excluding any signals below the detection limit, and setting this value to 100%. Each individual metabolite intensity was then calculated as a percentage of its respective sample sum, and these values were used for all fold-change and p-value calculations. The resulting sum-normalize blank corrected data matrix was then used for all group-wise comparisons, and t-tests were performed with the Python SciPy (1.1.0) (Jones et al., 2001) library to test for differences and generate statistics for downstream analyses. Any metabolite with p-value < 0.05 was considered significantly regulated (up or down). Heatmaps were generated with hierarchical clustering performed on the imputed matrix values utilizing the R library pheatmap (1.0.12) (Kolde and Kolde, 2015). In order to adjust for significant covariate effects (as applicable) in the experimental design the R package, DESeq2 (1.24.0) (Love et al., 2014) was used to test for significant differences. Data processing for this correction required the blank corrected matrix to be imputed with zeroes for non-detected values instead of the blank threshold to avoid false positives. This corrected matrix was then analyzed utilizing DESeq2 to calculate the adjusted p-value in the covariate model. Metabolic pathway analysis (integrating pathway enrichment analysis and pathway topology analysis) in Fig. S7F,G was performed using the Pathway Analysis module of MetaboAnalyst 5.0. Mus musculus (KEGG) was selected as the pathway library and hypergeometric test was the selected overrepresentation analysis method.

### Quantification and statistical analysis

All numerical results are presented as median with individual values shown in graphs. Normal distribution of data was not assumed, and statistical significance of differences between experimental groups was determined by nonparametric Mann–Whitney test. Differences were considered significant for P values <0.05 (*), <0.01 (**), and <0.001 (***). The number of animals per group (*n*) is indicated in the respective figure legends. All statistical calculations were performed using Prism (GraphPad) except for expression and chromatin analysis, for which R (v3.1.1 or v3.5.3) was used.

---

**Supplemental Table.**
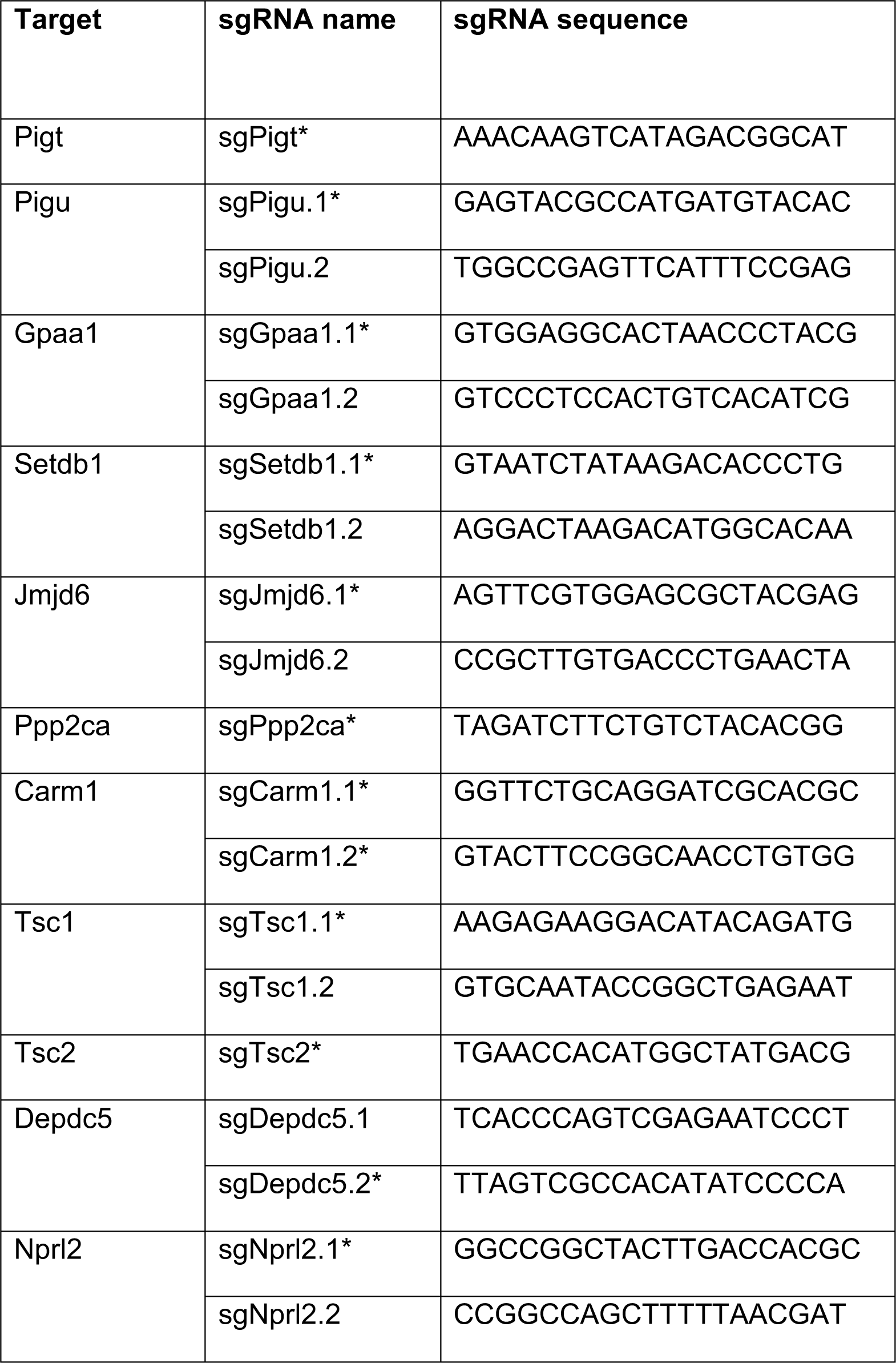
Sequences of sgRNAs used to target individual genes Asterisks denote sgRNAs used for experiments shown in Fig. 4-6

**Figure S1.**
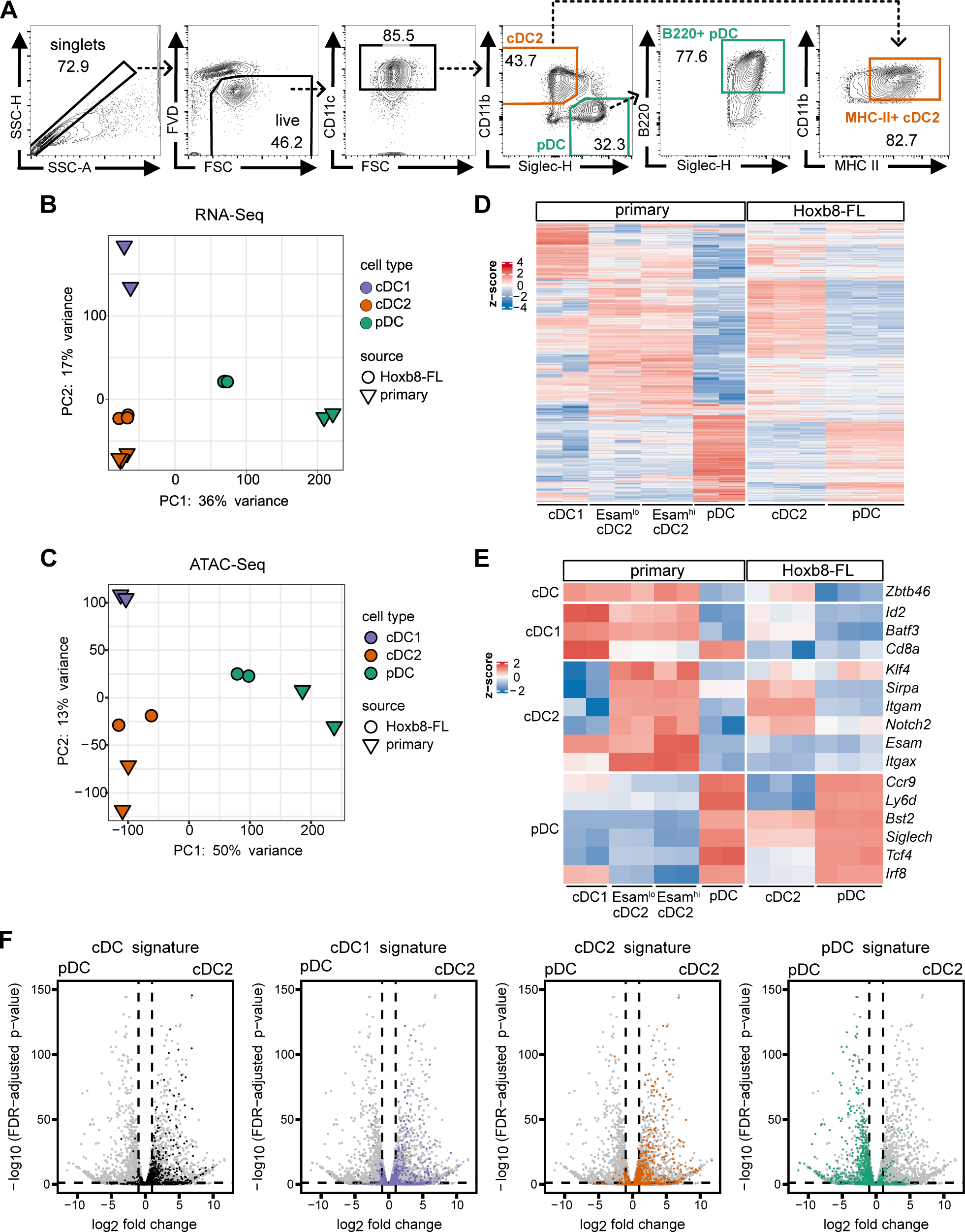
Additional characterization of Hoxb8-FL-derived DC subsets. (A) DC development in Hoxb8-FL cultures at day 7 of differentiation post removal of β- estradiol. Shown are representative staining plots highlighting SiglecH^+^ B220^+^ pDCs and CD11b^+^ MHCII^+^ cDC2. (B-C) Principal component analysis of cell source-corrected global RNA-Seq (panel B) and ATAC-Seq (panel C) profiles of primary splenic DCs (Lau et al., 2018) and Hoxb8- FL-derived DCs. (D) Gene expression in primary and Hoxb8-FL-derived DC subsets. Shown is the heatmap of 1,000 most variable genes (rows) across all cell types (columns, biological replicates); data represent cell source-corrected, row-scaled log2-transformed RNA-Seq expression values. (E) Select DC subset-specific genes from Fig. 1G. (F) Differential gene expression between Hoxb8-FL-derived cDC2 and pDC subsets. Shown are volcano plots (log2 fold change vs -log10 of false discovery rate-adjusted p-values) of gene expression with the signature genes of primary DC subsets (Lau et al., 2018) highlighted. Dashed lines mark p = 0.05 and log2 fold change of -1 and 1

**Figure S2.**
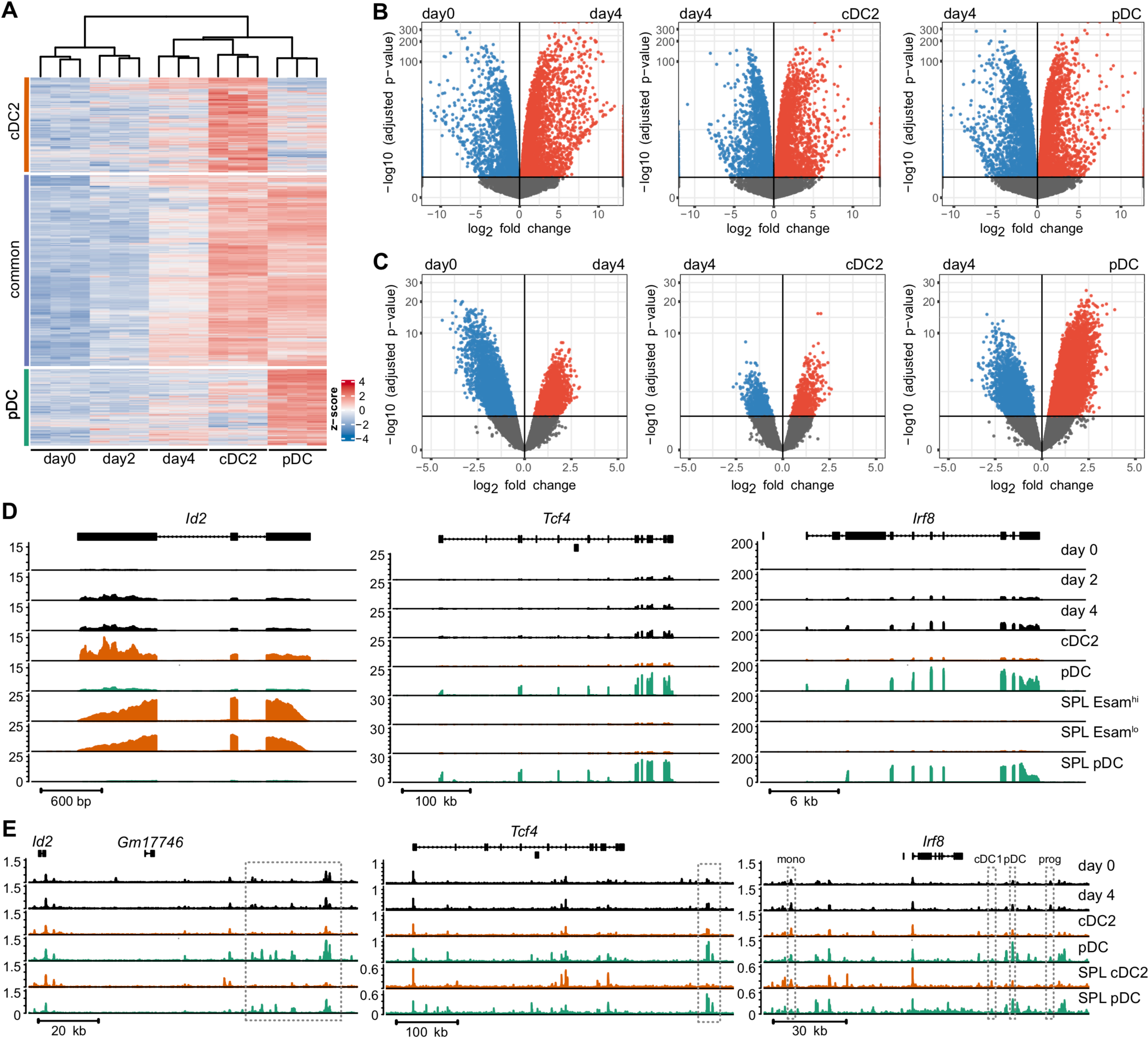
Characterization of DC differentiation from Hoxb8-FL progenitors. (A) Expression of transcripts upregulated in Hoxb8-FL-derived DCs during differentiation. Heatmap shows row-scaled log2-transformed gene expression values in biological replicates (columns) of Hoxb8-FL cells at the indicated days of differentiation, separately for cDC2- or pDC-enriched or common DC-enriched genes (adjusted p value < 0.05, log2 fold change > 1 compared to day 0). (B-C) Pairwise comparison of RNA-Seq (panel B) and ATAC-Seq (panel C) profiles of differentiating Hoxb8-FL cells on day 4 with undifferentiated progenitors (day 0) or differentiated cDC2s and pDCs. Shown are volcano plots (log2 fold change vs –log10 of adjusted p-values) of transcript or peak values. Significantly downregulated or upregulated features are indicated in blue and red, respectively; horizontal line marks adjusted p-value = 0.1 (D-E) Representative RNA-Seq (panel D) and ATAC-Seq (panel E) profiles of genes encoding DC-enriched transcription factors in differentiating Hoxb8-FL cells and primary splenic DCs. Shown are normalized read counts across the indicated genomic loci. Dashed squares indicate the negative distal regulatory region of *Id2* (Ghosh et al., 2014); pDC-specific downstream enhancer of *Tcf4* (Grajkowska et al., 2017); and enhancers of *Irf8* specific for pDCs, cDC1s, monocytes (mono) and DC progenitors (prog) (Bagadia et al., 2019; Murakami et al., 2021).

**Figure S3.**
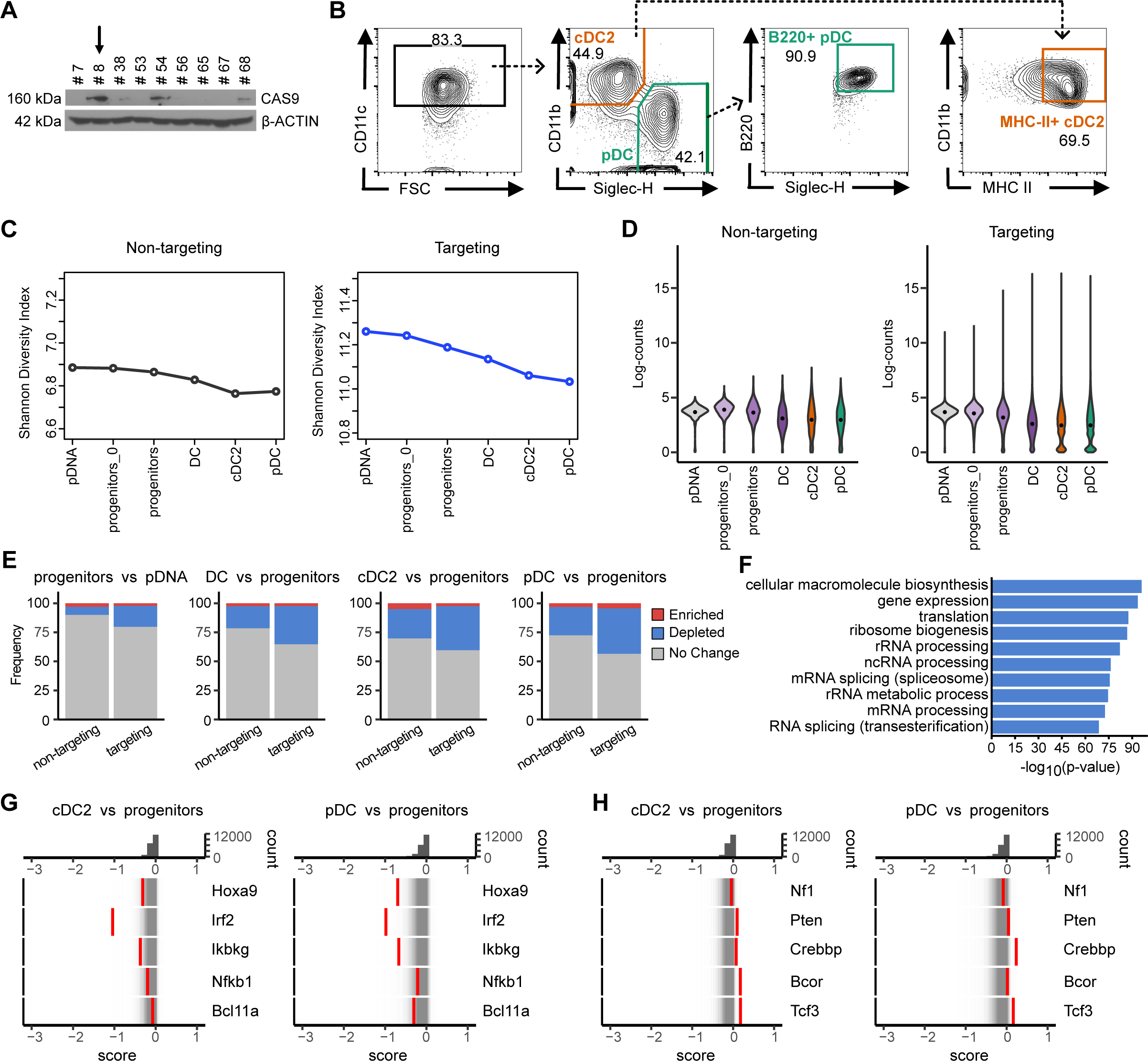
Additional characterization of genome-wide screen in Hoxb8-FL cells. (A) Western blot of Hoxb8-FL single cell clones transduced with a lentivirus expressing CAS9, probed for CAS9 protein or β-ACTIN as a housekeeping control. Arrow indicates the clone used for the CRISPR/Cas9 screen. (B) Differentiation capacity of the Cas9-expressing Hoxb8-FL cell clone indicated in panel A (gated on live cells). (C) Shannon diversity index of the control non-targeting sgRNAs (left panel) and targeting sgRNAs (right panel) in the samples shown in Fig. 3A. (D) Violin plots showing the distribution of log-transformed counts of the control non-targeting sgRNAs (left panel) and targeting sgRNAs (right panel) in the samples shown in Fig. 3A. (E) Fractions of significantly enriched (red) or depleted (blue) non-targeting or targeting sgRNAs (FDR <0.05) in pairwise comparisons between the indicated samples. (F) Enrichment of Gene Ontology (GO) biological processes in the genes targeted by significantly depleted guides in progenitors vs pDNA. Shown are bar charts of -log10 (p-value) of the top 10 enriched terms. (G-H) Rug plots of gene distribution between cDC2s or pDCs vs progenitors, as measured by the CRISPTimeR score. Top panels show the distribution of scores; bottom panels show individual genes (red) against the distribution of all genes (grey). (G) Rug plots of select genes showing sgRNA depletion in DCs. (H) Rug plots of select genes showing sgRNA enrichment in progenitors and/or DCs.

**Figure S4.**
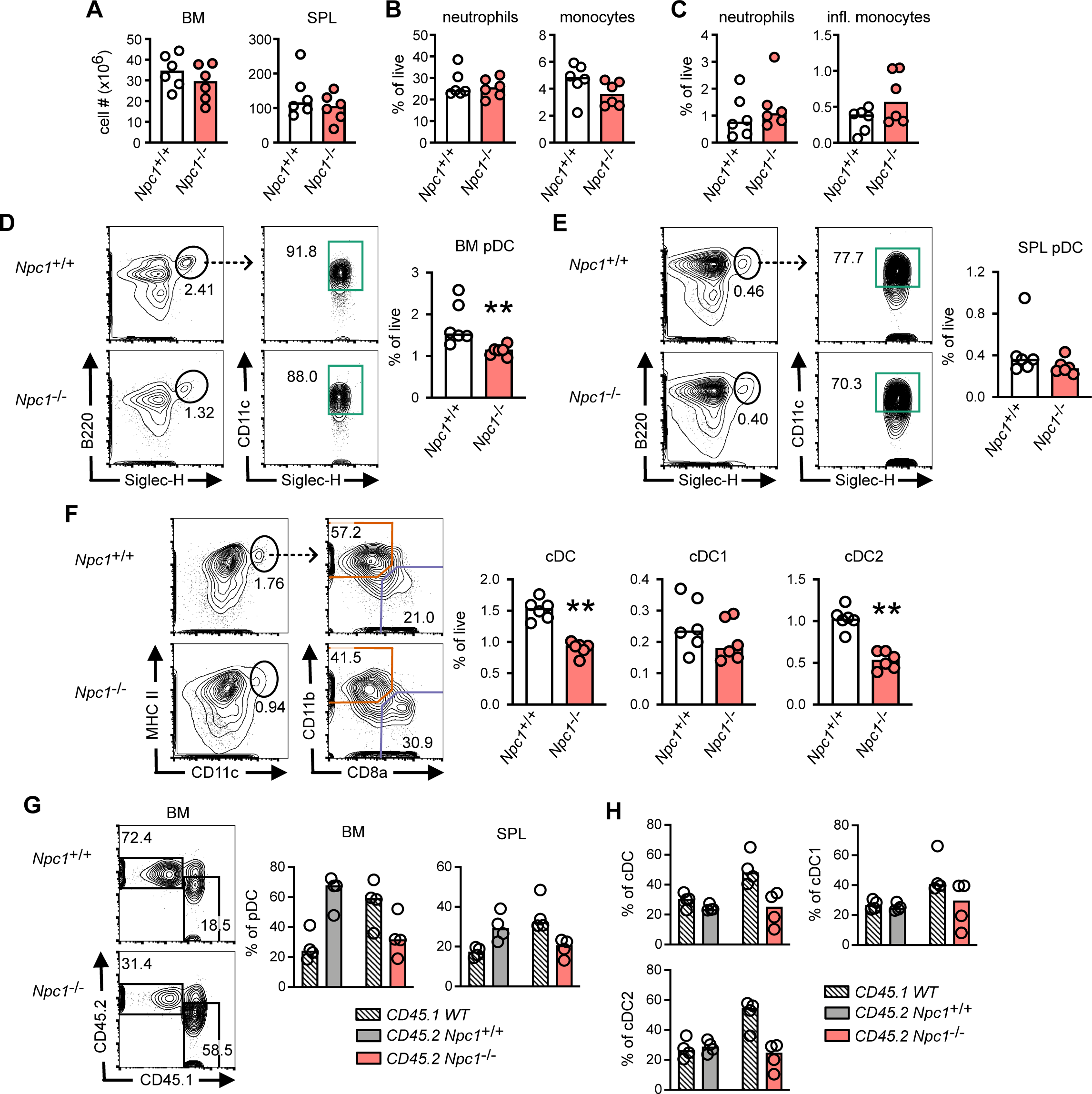
*Npc1* facilitates the development of specific DC subsets *in vivo*. (A-F) Analysis of young adult *Npc1*^-/-^ mice and control *Npc1*^+/+^ littermates. Symbols in bar graphs represent individual mice (n = 6) pooled from four experiments; bars represent the median. (A) Total cellularity in the BM and spleen. (B-C) The fraction of neutrophils (CD11b^+^ Ly6G^+^ Ly6C^lo^) and monocytes (CD11b^+^ Ly6G^-^ Ly6C^hi^) in BM (panel B) and spleen (panel C) as assessed by flow cytometry analysis. Shown are the frequencies among total live cells. (D-E) The fraction of pDCs in the BM (panel D) and spleen (panel E). Shown are representative staining profiles and the frequencies of B220^+^ SiglecH^+^ CD11c^+^ among total live cells. (F) The fraction of cDCs in the spleen. Shown are representative staining profiles and the frequencies of CD11c^hi^ MHCII^+^ cDCs, CD8a^+^ cDC1s and CD11b^+^ cDC2s among total live cells. (G-H) CD45.1^+^ recipient mice were reconstituted with CD45.2^+^ BM cells from *Npc1*^-/-^ or control *Npc1*^+/+^ mice mixed 1:1 with BM from CD45.1^+^ wild-type mice, and analyzed four months later. Symbols in bar graphs represent individual mice (n = 4); bars represent the median. (G) Donor-derived pDCs in the BM and spleen. Shown is representative staining of the gated pDCs in BM for donor-derived (CD45.2^+^) and competitor-derived (CD45.1^+^) cells and the frequencies of donor-derived and competitor-derived cells among pDCs in BM and spleen. (H) Donor-derived cDCs in the spleen. Shown are the frequencies of donor-derived and competitor-derived cells among splenic cDC subsets. Statistical significance was determined by Mann–Whitney test (**, *p* < 0.01).

**Figure S5.**
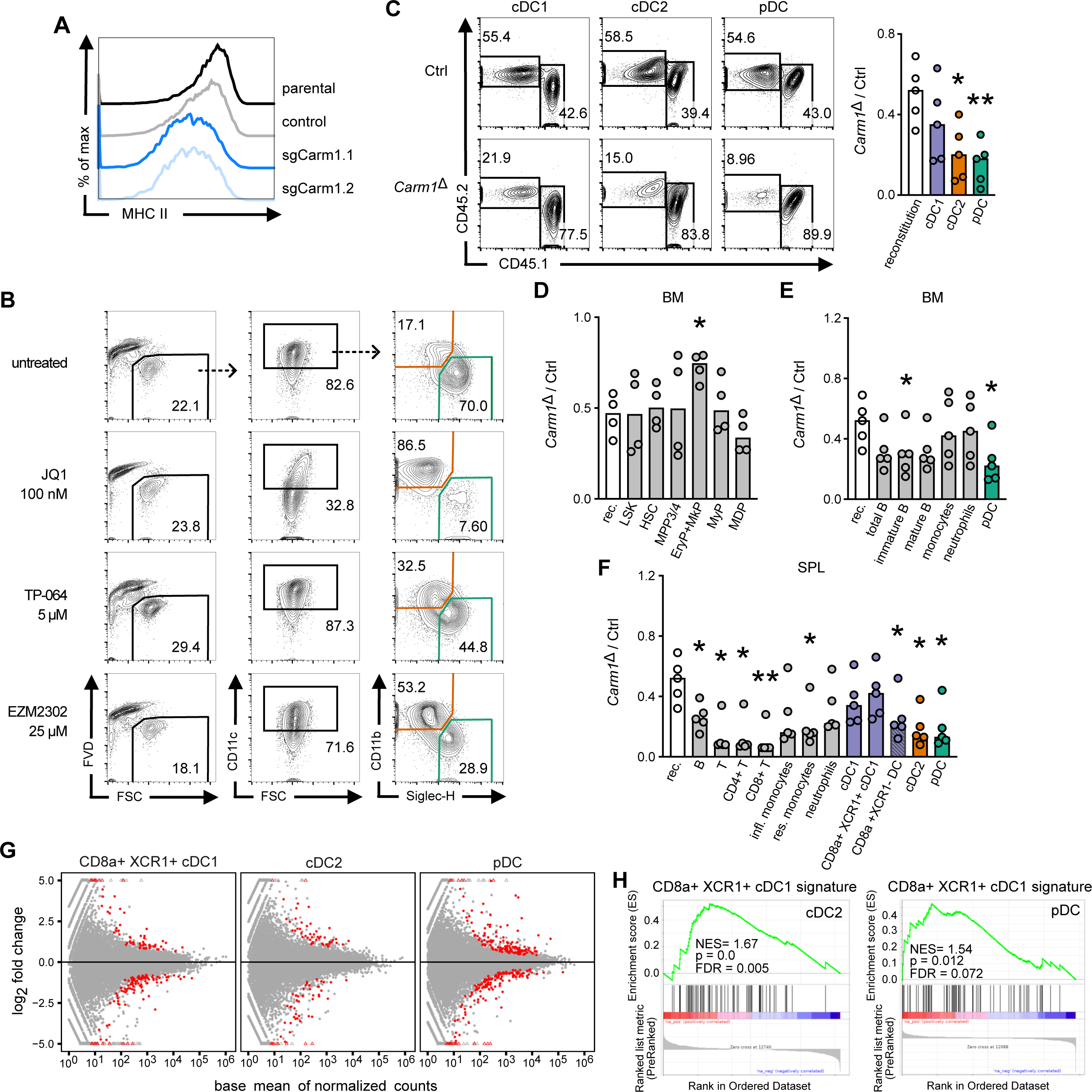
The role of *Carm1* in DC differentiation. (A) Expression of MHC cl II on gated CD11b^+^ cDC2s from parental, control or *Carm1*- deficient Hoxb8-FL cells described in Fig. 5A-B. (B) The effect of small molecule CARM1 inhibitors on the differentiation of Hoxb8-FL cells. Shown are staining profiles of Hoxb8-FL cells after 7 days of differentiation in the presence of CARM1 inhibitors TP-064 or EZM2302. The inhibitor of BRD proteins JQ1, which specifically impairs pDC development (Grajkowska et al., 2017), was used as a control. Representative of two experiments. (C-F) CD45.1^+^ recipient mice were reconstituted with BM cells from *Carm1*^fl/fl^ *Vav1*-Cre (designated *Carm1*^Δ^) mice or control (*Carm1*^fl/fl^) CD45.2^+^ mice mixed 1:1 with BM from CD45.1^+^ wild-type mice, and analyzed four months later. (C) DC development in Flt3L-supplemented BM cultures. Shown is representative staining profile of gated DC subsets for donor-derived (CD45.2^+^) and competitor-derived (CD45.1^+^) cells on day 7 cultures. Bar graphs show the ratio of *Carm1*^Δ^ to control donor-derived cells among DC subsets. Symbols represent cultures from individual recipient mice (n= 5); bars represent the median. (D-F) Ratios of *Carm1*^Δ^ to control CD45.2^+^ donor-derived cells among HSC (Lin^-^ ckit^+^ Sca1^+^ CD48^-^ CD150^+^) and progenitor cell populations (MPP3/4: Lin^-^ ckit^+^ Sca1^+^ CD48^+^ CD150^-^; EryP+MkP: Lin^-^ ckit^hi^ Sca1^-^ CD150^+^; MyP: Lin^-^ ckit^hi^ Sca1^-^ CD150^-^; MDP: Lin^-^ ckit^+^ Sca1^-^ Flt3^+^) in the BM (panel D) and mature cell types in the BM (panel E) and spleen (panel F). Lineage was defined as Ter119^+^ B220^+^ TCRβ^+^ CD11b^+^ Gr1^+^ NK1.1^+^. Symbols represent individual recipient mice (n=4-5); bars represent the median. (G-H) Transcriptome analysis of Carm1-deficient primary DCs. Primary splenic DC subsets (XCR1+ cDC1, cDC2, pDC) were sorted from 3 individual chimeric mice reconstituted with BM from *Carm1*^Δ^ or control donors and analyzed by RNA-seq. (G) Pairwise comparisons of expression profiles within each DC subset (average normalized counts across all samples vs moderated log2 fold change of *Carm1*^Δ^ over control); genes with significant differential expression (adjusted p value < 0.05) are highlighted in red. (H) Gene Set Enrichment Analysis (GSEA) of RNA-Seq data testing the enrichment of the cDC1 transcriptional signature in *Carm1*^Δ^ vs control cDC2s and pDCs. Statistical significance by Mann-Whitney test (*, P < 0.05; **, P < 0.01)

**Figure S6.**
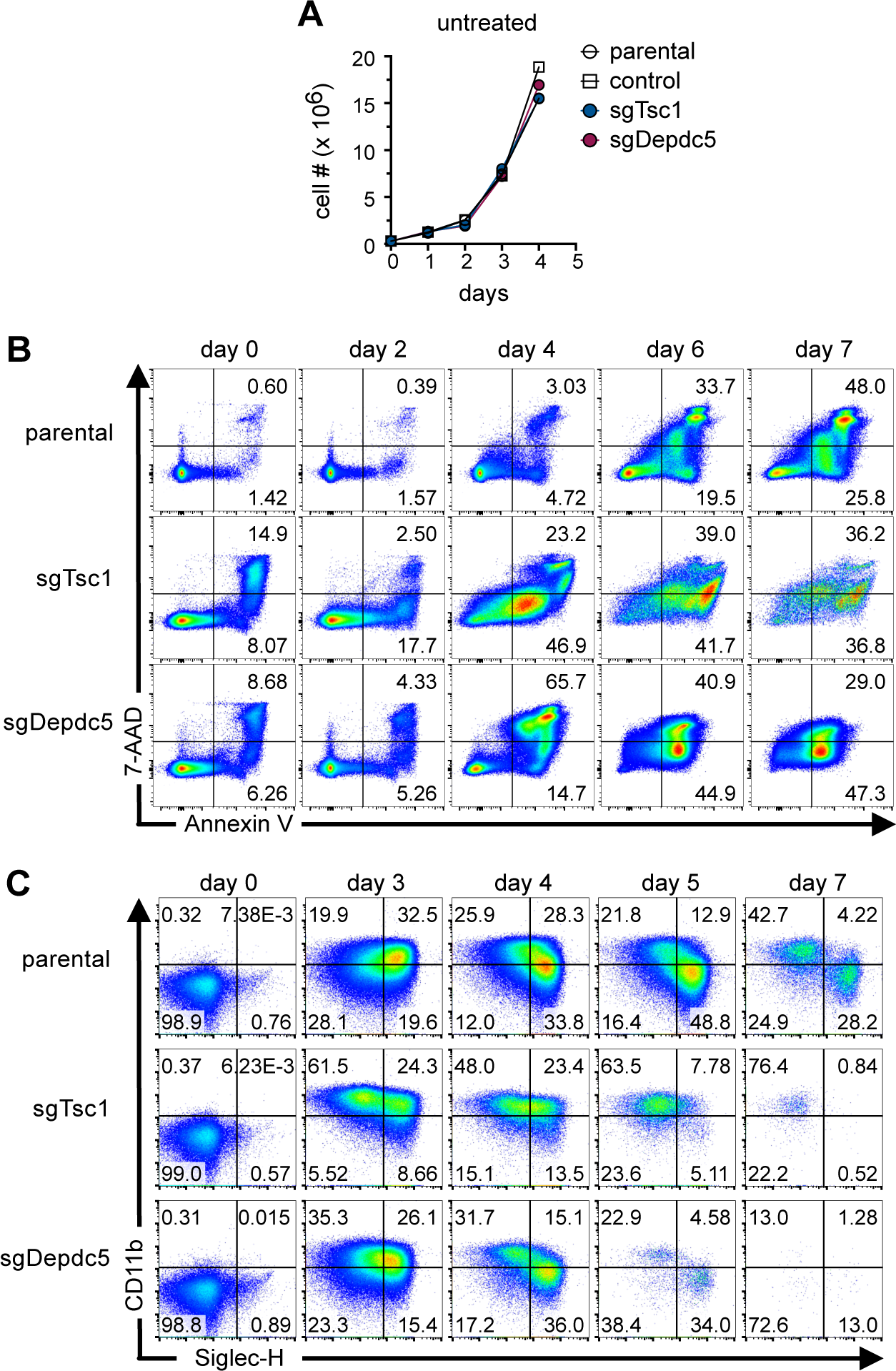
Characterization of Hoxb8-FL cells with targeted TSC or GATOR1 complexes. Hoxb8-FL cells were transduced with control sgRNA or sgRNAs targeting *Tsc1* or *Depdc5,* encoding subunits of the respective TSC and GATOR1 complexes. (A) Growth curves of parental, control or sgRNA-transduced Hoxb8-FL cells as progenitors in undifferentiated conditions. (B) Apoptosis of parental or sgRNA-transduced Hoxb8-FL cells during differentiation. Total cells were stained to detect apoptosis (Annexin V) and membrane permeability (7- AAD) at the indicated days of DC differentiation. Cells were gated on total single cells. (C) The phenotype of parental or sgRNA-transduced Hoxb8-FL cells at the indicated days of DC differentiation. Cells were gated on viable CD11c^+^ singlets.

**Figure S7.**
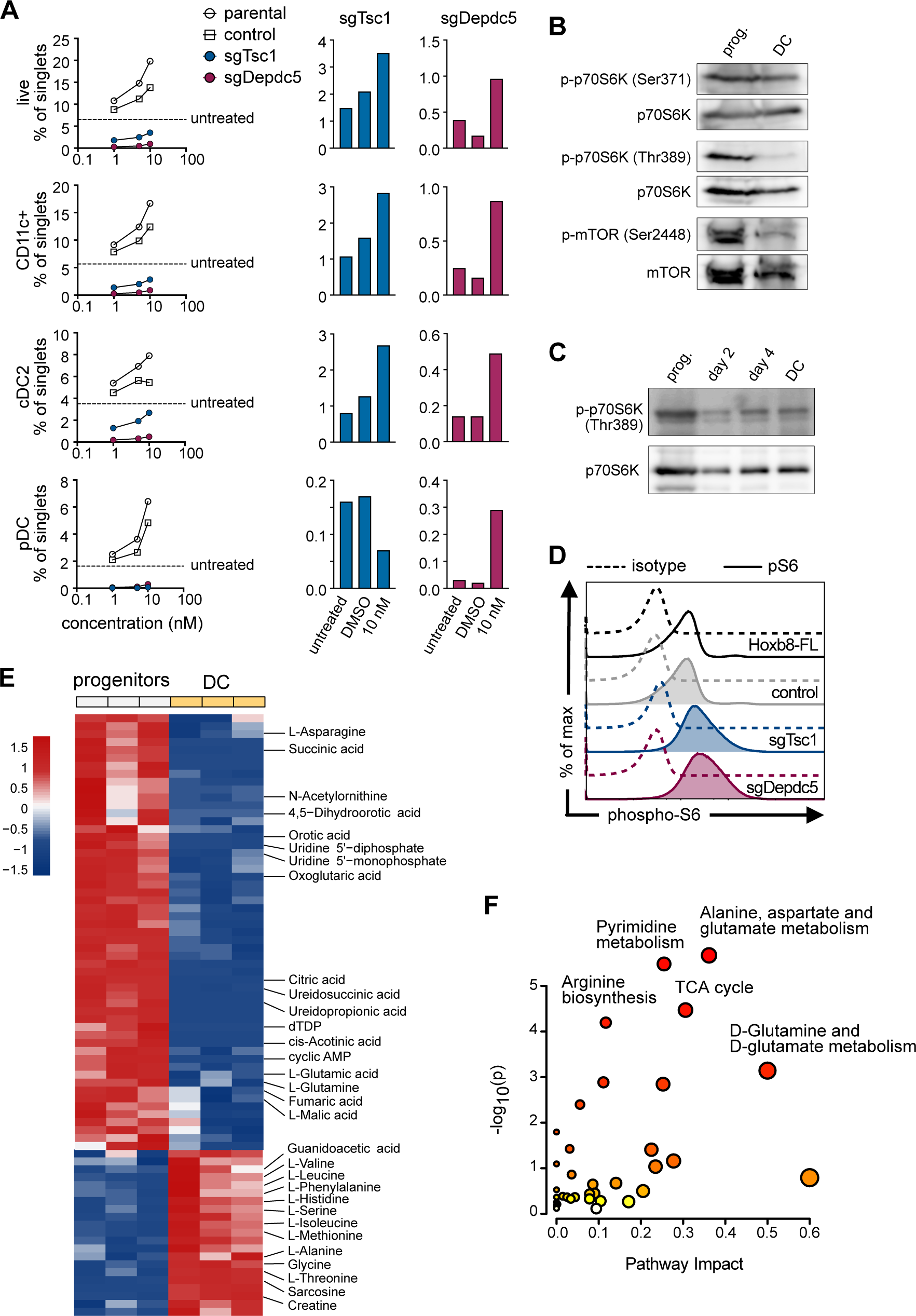
Characterization of mTOR signaling during DC differentiation of Hoxb8- FL cells. (A) Differentiation of Hoxb8-FL cells with targeted components of TSC and GATOR1 complexes in the presence of mTOR inhibitor. Parental Hoxb8-FL cells or cells transduced with control sgRNA or sgRNAs targeting indicated genes encoding TSC or GATOR1 subunits were differentiated into DCs in the presence of Torin1 and analyzed by flow cytometry. Shown are frequencies of live cells, CD11c+ DCs and DC subsets among single cells at the indicated concentrations of Torin1. Bar graphs indicate the frequencies of corresponding cell subsets in the presence of 10 nM Torin1 compared to untreated or vehicle only controls. (B-C) Western blot of mTOR pathway proteins in Hoxb8-FL cells. (B) Protein lysates from Hoxb8-FL undifferentiated progenitors (prog.) or bulk DCs on day 7 of differentiation were probed for p70S6K phosphorylated at different residues (Ser371 or Thr389) and for mTOR phosphorylated at Ser2448, along with total p70S6K and mTOR. (C) Protein lysates from Hoxb8-FL undifferentiated progenitors (prog.) or bulk cells at the indicated days of differentiation were probed for p70 S6K phosphorylated at Thr389 along with total p70S6K. (D) Intracellular staining for phospho-S6 in live undifferentiated parental Hoxb8-FL cells, cells transduced with control sgRNA or sgRNAs targeting indicated genes encoding TSC or GATOR1 subunits. Dashed histograms indicate isotype controls. Representative staining profiles of two experiments are shown. (E) Differentially represented metabolites in Hoxb8-FL progenitors and Hoxb8-FL-derived DCs. Heatmap shows relative abundance of metabolites (rows; row-scaled z-score transformed) across the two cell types; columns represent biological replicates. (F) Pathway analysis of metabolites enriched in Hoxb8-FL progenitors identified in panel E. Metabolic pathways with p < 0.001 as determined by enrichment analysis are annotated.

